# Modeling mTORopathy-related epilepsy in cultured murine hippocampal neurons using the multi-electrode array

**DOI:** 10.1101/2024.04.29.591584

**Authors:** Anouk M. Heuvelmans, Martina Proietti Onori, Monica Frega, Jeffrey D. de Hoogen, Eveline Nel, Ype Elgersma, Geeske M. van Woerden

## Abstract

The mechanistic target of rapamycin complex 1 (mTORC1) signaling pathway is a ubiquitous cellular pathway. mTORopathies, a group of disorders characterized by hyperactivity of the mTORC1 pathway, illustrate the prominent role of the mTOR pathway in disease pathology, often profoundly affecting the central nervous system. One of the most debilitating symptoms of mTORopathies is drug-resistant epilepsy, emphasizing the urgent need for a deeper understanding of disease mechanisms to develop novel anti-epileptic drugs. In this study, we explored the multiwell Multi-electrode array (MEA) system as a tool to identify robust network activity parameters in an approach to model mTORopathy-related epilepsy *in vitro*. To this extent, we cultured mouse primary hippocampal neurons on the multiwell MEA to identify robust network activity phenotypes in mTORC1-hyperactive neuronal networks. mTOR-hyperactivity was induced either through deletion of *Tsc1* or overexpression of a constitutively active RHEB variant identified in patients, RHEBp.P37L. mTORC1 dependency of the phenotypes was assessed using rapamycin, and vigabatrin was applied to treat epilepsy-like phenotypes. We show that hyperactivity of the mTORC1 pathway leads to aberrant network activity. In both the *Tsc1*-KO and RHEB-p.P37L models, we identified changes in network synchronicity, rhythmicity, and burst characteristics. The presence of these phenotypes is prevented upon early treatment with the mTORC1-inhibitor rapamycin. Application of rapamycin in mature neuronal cultures could only partially rescue the network activity phenotypes. Additionally, treatment with the anti-epileptic drug vigabatrin reduced network activity and restored burst characteristics. Taken together, we showed that mTORC1-hyperactive neuronal cultures on the multiwell MEA system present reliable network activity phenotypes that can be used as an assay to explore the potency of new drug treatments targeting epilepsy in mTORopathy patients and may give more insights into the pathophysiological mechanisms underlying epilepsy in these patients.

**ABBREVIATIONS**

AED, anti-epileptic drug, CoV^NIBI^, coefficient of variance of NIBI, CTR, control transduced cultures, DIV, days in vitro, DMEM, Dulbecco’s modified Eagle medium, DMSO, dimethyl sulfoxide, GABA, gamma-aminobutyric acidergic, GAPDH, Glyceraldehyde-3-Phosphate Dehydrogenase, iPSC, induced pluripotent stem cell, KO, knock-out, LV, lentivirus, MEA, multi-electrode array, MFR, mean firing rate, mTORC1, mechanistic target of rapamycin complex 1, NB, network burst, NBC, network burst composition, NBD, network burst duration, NBM, neurobasal medium, NBR, network burst rate, NIBI, network interburst interval, NT, non-transduced, RHEB, Ras-homolog enriched in brain, %RS, percentage of random spikes, TBS, tris buffered saline, TSC, Tuberous sclerosis complex, WT, wildtype

## 1. INTRODUCTION

The mTORC1 (mammalian target of rapamycin complex 1) pathway is a ubiquitous cellular pathway. It integrates signals from the intra-and extracellular environment to regulate fundamental cellular processes such as protein synthesis, metabolism, and autophagy (1). Studies on the function of mTORC1 revealed it plays an important role in neurodevelopment, as it was found to be involved in the regulation of neural differentiation, migration and survival, neuronal morphology, membrane excitability, and synaptic transmission (2–4). Based on this, it is not surprising that hyperactivity of the mTORC1 pathway has been observed in multiple neurodevelopmental disorders, collectively referred to as mTORopathies. Shared pathological hallmarks of these disorders are hemimegalencephaly, focal cortical dysplasia, gliosis, the presence of dysmorphic neurons, and epilepsy (5).

Many of the mTORopathies are caused by pathogenic variants in genes encoding proteins of the mTORC1 signaling pathway, resulting in hyperactivity of the pathway (6,7). The most prevalent and well-studied example of an mTORopathy is Tuberous Sclerosis Complex (TSC). TSC patients carry a loss of function (LoF) mutation in either the *TSC1* or *TSC2* gene, resulting in loss of the encoded proteins Hamartin and Tuberin respectively. These proteins form a complex that inhibits Ras-Homolog Enriched in Brain (RHEB), by converting it from an active GTP-bound to an inactive GDP-bound state. In turn, RHEB is a direct activator of the mTORC1 complex. Loss of a functional TSC complex thus leads to increased RHEB activity resulting and concomitant mTORC1 hyperactivity. More recently, gain of function mutations in *RHEB* have been identified that directly cause mTORC1 hyperactivity (8).

Many mTORopathy patients suffer from severe and often intractable epilepsy, which is reported as one of the most debilitating symptoms of the disorder (5,9). Moreover, intractable epilepsy is strongly associated with intellectual disability and autism. Patients generally respond poorly to anti-epileptic drugs (AED) and a large cohort study with TSC patients showed only a few patients remain seizure-free for more than a year (10). Rapamycin, an allosteric mTORC1 inhibitor, has proven to be effective in reducing mTORC1 hyperactivity in the brain (11). Although Rapamycin and its analogs have been shown to reduce seizure frequency and severity, patients are not seizure-free and often suffer from serious side effects (12). This emphasizes the urgent need for a better understanding of disease mechanisms to allow the development of novel anti-epileptic drugs.

Epilepsy has been studied *in vitro* using multi-electrode arrays (MEAs) (13–17). MEAs can be used to record network activity of cultured neurons over weeks, even up to months. During the first weeks in culture, neuronal activity develops from random spiking into robust network activity characterized by rhythmic and synchronous bursting (18,19). Several previous experiments have used MEAs to model TSC. A study by Bateup reported an increased firing rate as a result of *Tsc1* knock-out in mouse hippocampal cultures, which was rescued by rapamycin treatment (20). Similarly, neurons derived from TSC1-and TSC2-patient induced pluripotent stem cells (iPSC) displayed significantly more spiking activity and increased synchronicity compared to control cultures (21–23). The MEA system holds the potential to provide a semi high-throughput screening method for potential new treatments. However, current studies have mostly focused on the readout parameters firing rate and network synchronicity, which were reported to be more prone to variation (24). Parameters shown to be more stable in MEA measurements are parameters such as frequency and duration of network bursts (24).

The current study explores the multiwell MEA system as a tool to identify changes in the more stable network activity parameters as an approach to study mTORopathy-related epilepsy *in vitro*. To this end, we made use of 2 different mTORopathy models in mouse primary hippocampal neurons: 1) deletion of *Tsc1* and 2) overexpression of a constitutively active RHEB variant identified in patients, RHEB-p.P37L (25,26), both shown to cause mTORC1 hyperactivity. We examined changes in network activity in response to mTORC1 hyperactivity and identified phenotypes that can serve as a proxy for epilepsy in these cultured neurons. This was substantiated through application of vigabatrin, the first-line anti-epileptic drug (AED) used in TSC patients (10), and previously shown to have the highest anti-epileptic effect in a *Tsc1-*KO mouse model (27). The identification of changes in network activity in this system, and their sensitivity to drug manipulations will allow us to perform more mechanistic studies to further disentangle the pathophysiology underlying mTORC1 hyperactivity-related epilepsies as well as to identify novel AEDs.

## 2. RESULTS

### 2.1 mTORC1 hyperactive cultures show alterations in network synchronicity and network rhythmicity

To induce mTORC1 hyperactivity, we made use of two different models. Mouse primary hippocampal neurons were cultured either from inducible *Tsc^f/f^* (27) or from FvB/NHsD wild-type mice. mTORC1 hyperactivity was induced through viral expression of Cre recombinase in the *Tsc^f/f^* neurons, resulting in homozygous knock-out of *Tsc1* (*Tsc1*-KO), or viral expression of the constitutively active RHEB variant RHEB-p.P37L in the FvB/NHsD hippocampal cultures (Fig 1A). We used phosphorylation of S6 (pS6^Ser240/244^), a specific substrate of mTORC1, as a read-out for mTORC1 activity to confirm hyperactivity of the pathway. As expected, expression of RHEB-p.P37L or deletion of *Tsc1* in mouse hippocampal cultures resulted in an increase of the pS6:S6 ratio (Fig 1B: *Tsc1*-KO: t(12) = 12.28, p <0.0001, RHEB-p.P37L: t(12) = 3.731, p = 0.0144), and of pS6 (Fig 1C: *Tsc1*-KO: t(12) = 27.97, p <0.0001, RHEB-p.P37L: t(12) = 8.192, p <0.0001), compared to control cultures. In previous studies it was already shown that overexpression of RHEB-WT increases mTORC1-hyperactivity, but to a lesser extent than the RHEB-p.P37L variant (25). Additionally, in contrast to RHEB-WT, this variant is resistant to inhibition by the TSC-complex (25,26). Treatment of the cultures with rapamycin, an allosteric mTORC1 inhibitor, decreased the pS6:S6 ratio compared to the non-treated control (Fig 1B: Control + rapamycin: t(12) = 8.233, p <0.0001, *Tsc1*-KO + rapamycin: t(12) = 7.915, p <0.0001, RHEB-p.P37L + rapamycin: t(12) = 7.382, p < 0.0001). Additionally, we confirmed that *Tsc1* was successfully deleted in our *Tsc1*^f/f^-cultures through the expression of Cre recombinase, as TSC1 (Hamartin) protein levels were decreased in the cultures transduced with the Cre expressing lentivirus (Fig 1D: t(4) = 15.23, p = 0.0001).

**Figure 1.**
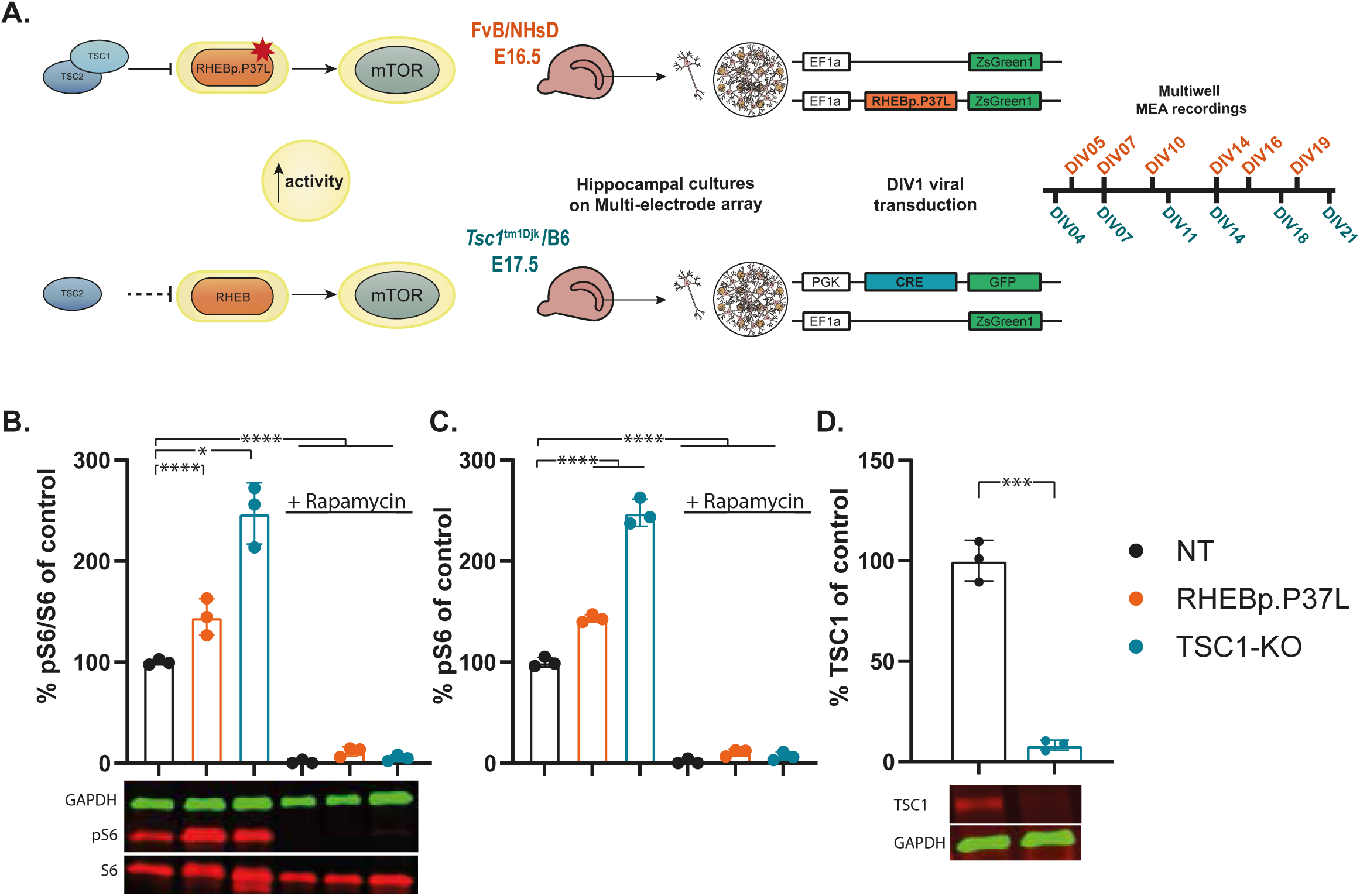
Upregulation of mTORC1 activity through viral transduction of neuronal cultures. (A) Experimental approach: mTOR hyperactivity is induced through transduction of FvB/NHsD neuronal cultures with a lentivirus expressing the gain-of-function RHEB-p.P37L variant (top illustration), or *Tsc1*^f/f^-cultures with a lentivirus expressing Cre-recombinase to induce *Tsc1*-KO (bottom illustration) at DIV01. Timeline of MEA recordings for each cell line. (B) pS6^(Ser240/244)^:S6 ratio in RHEB-p.P37L and *Tsc1*-KO cultures are increased compared to control. Rapamycin treatment reduces pS6^(Ser240/244)^:S6 ratio compared to control (N = 3 wells / group). (C) pS6^(Ser240/244)^ levels in RHEB-p.P37L and *Tsc1*-KO cultures are increased compared to control. Rapamycin treatment reduces pS6^(Ser240/244)^ levels compared to control (N = 3 wells / group). (D) Reduced TSC1 levels in transduced *Tsc1*^f/f^ cultures compared to control (N = 3 wells/group). Western blot data are presented as mean ± SD. Significant differences in the pS6:S6 ratio were determined using a one-way ANOVA with post hoc Bonferroni correction. TSC1 levels were compared using an unpaired Student’s t-test. *p < 0.05, **p < 0.01, ***p < 0.001, ****p < 0.0001.

We recorded the activity of cultures using the multiwell MEA system across 3 weeks in vitro (Fig 1A), during which the neuronal activity pattern developed from random spikes to synchronized bursting activity (Fig 2A-B) (18,19). In these electrophysiological experiments, we controlled for the increased RHEB levels, by taking along lentiviral expression of the RHEB-WT (Fig S1). In the RHEB-p.P37L cultures, we found an increase in mean firing rate (MFR), starting from DIV07 (Fig 2C_1_). In the *Tsc1*-KO cultures, this phenotype appeared from DIV11 (Fig 2C_2_). An earlier phenotype in the RHEB-p.P37L culture was expected since the RHEB-p.P37L protein is expressed soon after lentiviral transduction, immediately inducing mTORC1 hyperactivity. In *Tsc1*-KO cultures, upregulation of mTORC1 activity is expected to take longer due to the Tsc1 protein half-life of 22 hours (28). After 1 week *in vitro*, neuronal cultures started to synchronize their activity into network-wide bursting events. These synchronous events increased during the second week *in vitro,* while non-synchronous, random activity decreased (18,19,29). We used the network burst rate (NBR, Fig 2D) and percentage of random spikes (%RS, Fig 2E), defined as the percentage of spikes that are not part of a network burst, as measures for network synchronicity. In control cultures, the network burst rate increased in the first two weeks after which it stabilized. From this time point onwards, cultures showed highly synchronized activity as most spikes clustered together in network bursts. Interestingly, synchronization of the network in mTORC1 hyperactive cultures was already observed earlier, as networks appeared more synchronous, already during the second week *in vitro* compared to control cultures (Fig 2D-E). The RHEB-p.P37L cultures showed an increased NBR at DIV07, and a reduced %RS compared to control at DIV07 and DIV11 (Fig 2D_1_-E_1_). Importantly, these effects of upregulation of the mTORC1 pathway through the expression of RHEB-p.P37L were mostly specific for the pathogenic variant, as in cultures expressing wild-type RHEB (RHEB-WT) the firing rate and network synchronicity were similar to control cultures (Fig S1A-C). In the *Tsc1*-KO cultures, NBR was increased from DIV11-18, and we detected a slight reduction of the %RS at DIV11 and DIV14 compared to controls (Fig 2D_2_-E_2_). All statistical comparisons are presented in Tables 1A-B and S1A-B.

**Figure 2.**
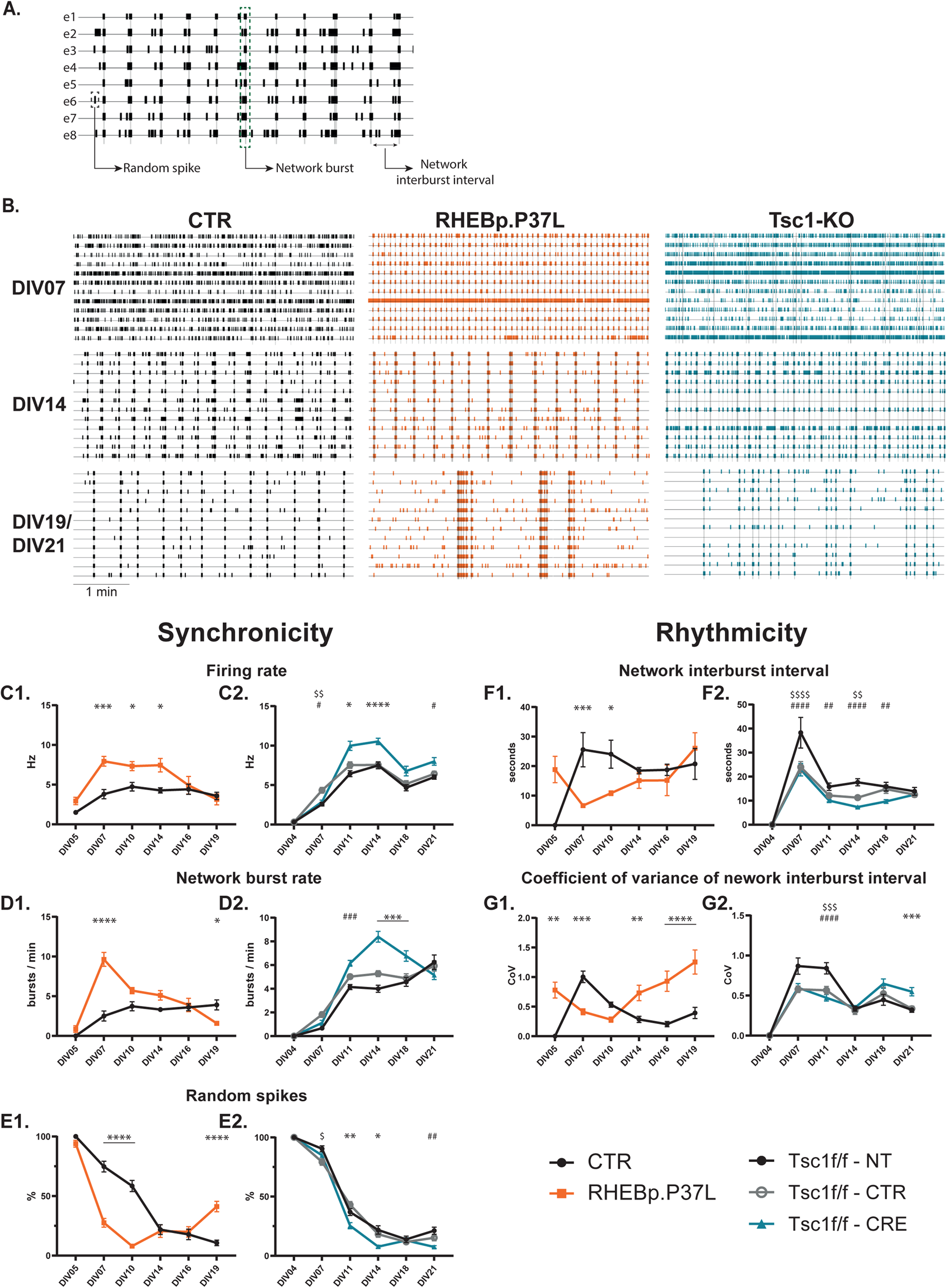
mTORC1 hyperactivity causes alterations in synchronicity and rhythmicity of neuronal networks on the MEA. (A) Example of an 8-electrode raster plot with network outcome parameters. Each horizontal line represents a single electrode, with vertical lines representing spikes. Gray bars spanning the entire raster represent network bursts. (B) Representative raster plots of 5-minute recordings in a single MEA well for control, RHEB-p.P37L, and *Tsc1*-KO cultures at 3 different time points (DIV07, DIV14, and DIV19 (CTR & RHEB-p.P37L) / DIV21 (*Tsc1*-KO)). (C-G) Quantification of network activity parameters in RHEB-p.P37L and *Tsc1*-KO cultures compared to their controls: Firing rate in Hz (C1-2), network burst rate in network bursts/min (D1-2), percentage random spikes (E1-2), network inter-burst interval (NIBI) in seconds (F1-2), and coefficient of variance of NIBI (G1-2). Data is presented as mean ± SEM. Sample size (wells/replicates): CTR: n = 15 / 3, RHEB-p.P37L: n = 17 / 3. *Tsc1*^f/f^-NT: n = 42 / 4, *Tsc1*^f/f^-CTR: n = 46 / 4, *Tsc1*^f/f^-CRE: n = 51 / 4. Significant differences were determined using a Mixed-effect analysis with post hoc Bonferroni correction. *p < 0.05, **p < 0.01, ***p < 0.001, ****p < 0.0001. (panels 2) * represent significant differences between *Tsc1*^f/f^-CRE and both controls, # represent significant differences between *Tsc1*^f/f^-CRE and either of the controls, $ represent significant differences between *Tsc1*^f/f^-NT and *Tsc1*^f/f^-CTR. All statistical comparisons are presented in Table 1A-B.

**Table 1A.**
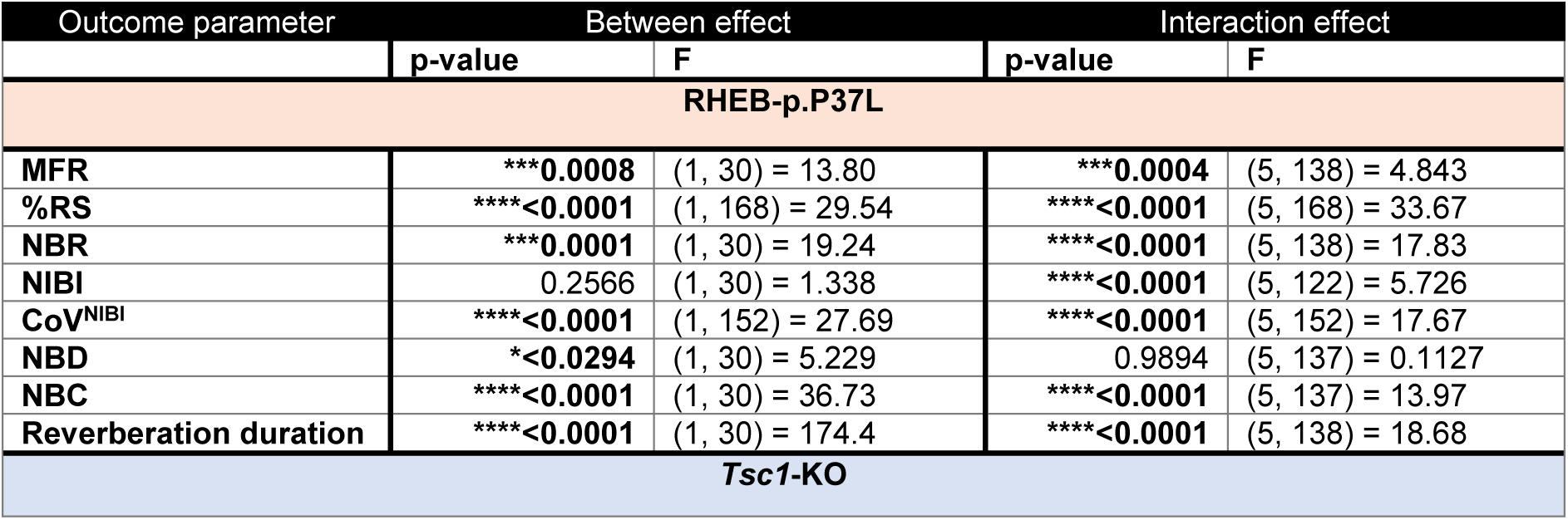
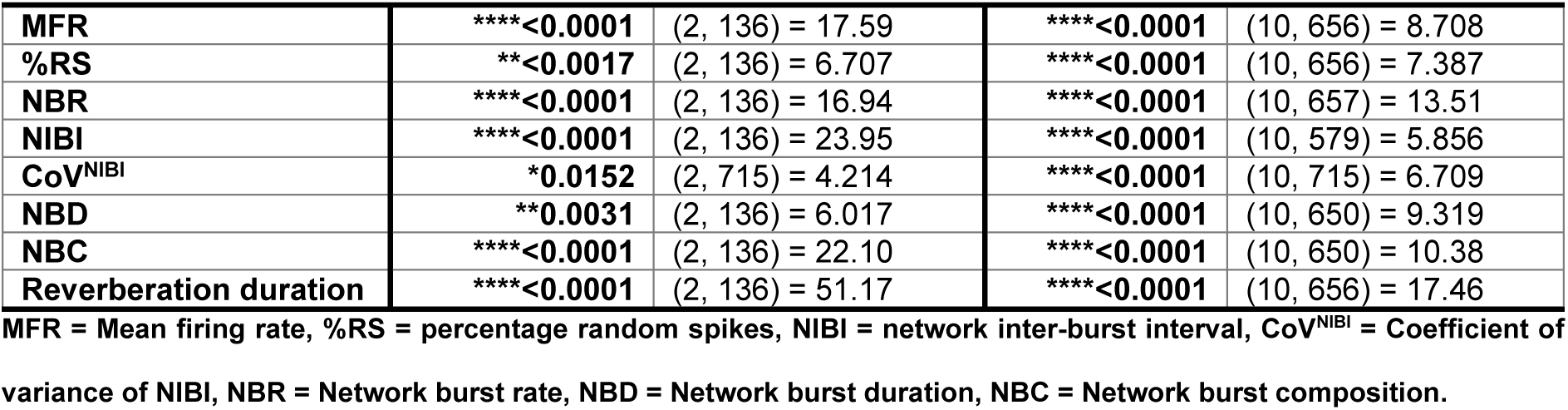
Statistical analysis overview of mixed effects analysis for establishment of mTORC1 hyperactivity-induced phenotypes on the MEA.

**Table 1B.** Post-hoc Bonferroni corrected multiple comparisons for establishment of mTORC1 hyperactivity-induced phenotypes on the MEA.

| Parameter | Comparison | p-value |  |  |  |  |  |
| --- | --- | --- | --- | --- | --- | --- | --- |
|  |  | DIV04/05 | DIV07 | DIV10/11 | DIV14 | DIV16/18 | DIV19/21 |
| MFR | CTR - RHEB-p.P37L | 0.0681 | **0.0002** | **0.0148** | **0.0132** | >0.9999 | >0.9999 |
|  | *Tsc1<sup>fl/fl</sup>*-NT - *Tsc1<sup>fl/fl</sup>*-CTR | >0.9999 | **0.0018** | 0.9513 | >0.9999 | >0.9999 | >0.9999 |
|  | *Tsc1<sup>fl/fl</sup>*-NT - *Tsc1<sup>fl/fl</sup>*-CRE | >0.9999 | >0.9999 | **<0.0001** | **<0.0001** | 0.0834 | **0.0095** |
|  | *Tsc1<sup>fl/fl</sup>*-CTR - *Tsc1<sup>fl/fl</sup>*-CRE | >0.9999 | **0.0146** | **0.0159** | **<0.0001** | 0.5289 | 0.1628 |
| %RS | CTR - RHEB-p.P37L | >0.9999 | **<0.0001** | **<0.0001** | >0.9999 | >0.9999 | **<0.0001** |
|  | *Tsc1<sup>fl/fl</sup>*-NT - *Tsc1<sup>fl/fl</sup>*-CTR | >0.9999 | **0.0113** | >0.9999 | >0.9999 | >0.9999 | >0.9999 |
|  | *Tsc1<sup>fl/fl</sup>*-NT - *Tsc1<sup>fl/fl</sup>*-CRE | >0.9999 | >0.9999 | **0.0053** | **0.0002** | >0.9999 | **0.0003** |
|  | *Tsc1<sup>fl/fl</sup>*-CTR - *Tsc1<sup>fl/fl</sup>*-CRE | >0.9999 | >0.9999 | **<0.0001** | **0.0131** | >0.9999 | 0.2011 |
| NBR | CTR - RHEB-p.P37L | >0.9999 | **<0.0001** | 0.0534 | 0.1059 | >0.9999 | **0.0270** |
|  | *Tsc1<sup>fl/fl</sup>*-NT - *Tsc1<sup>fl/fl</sup>*-CTR | >0.9999 | 0.1970 | 0.9145 | 0.0789 | >0.9999 | >0.9999 |
|  | *Tsc1<sup>fl/fl</sup>*-NT - *Tsc1<sup>fl/fl</sup>*-CRE | >0.9999 | >0.9999 | **0.0001** | **<0.0001** | **<0.0001** | 0.2384 |
|  | *Tsc1<sup>fl/fl</sup>*-CTR - *Tsc1<sup>fl/fl</sup>*-CRE | >0.9999 | >0.9999 | 0.1552 | **<0.0001** | **0.0007** | >0.9999 |
| NIBI | CTR - RHEB-p.P37L | 0.0828 | **0.0006** | **0.0264** | >0.9999 | >0.9999 | >0.9999 |
|  | *Tsc1<sup>fl/fl</sup>*-NT - *Tsc1<sup>fl/fl</sup>*-CTR | >0.9999 | **<0.0001** | 0.5094 | **0.0023** | >0.9999 | >0.9999 |
|  | *Tsc1<sup>fl/fl</sup>*-NT - *Tsc1<sup>fl/fl</sup>*-CRE | >0.9999 | **<0.0001** | **0.0069** | **<0.0001** | **0.0046** | >0.9999 |
|  | *Tsc1<sup>fl/fl</sup>*-CTR - *Tsc1<sup>fl/fl</sup>*-CRE | >0.9999 | >0.9999 | >0.9999 | 0.2361 | 0.0574 | >0.9999 |
| CoV<sup>NIBI</sup> | CTR - RHEB-p.P37L | **0.0016** | **0.0003** | 0.3313 | **0.0050** | **<0.0001** | **<0.0001** |
|  | *Tsc1<sup>fl/fl</sup>*-NT - *Tsc1<sup>fl/fl</sup>*-CTR | >0.9999 | 0.0678 | **0.0003** | >0.9999 | >0.9999 | >0.9999 |
|  | *Tsc1<sup>fl/fl</sup>*-NT - *Tsc1<sup>fl/fl</sup>*-CRE | >0.9999 | 0.2038 | **<0.0001** | >0.9999 | 0.0615 | **0.0050** |
|  | *Tsc1<sup>fl/fl</sup>*-CTR - *Tsc1<sup>fl/fl</sup>*-CRE | >0.9999 | >0.9999 | >0.9999 | >0.9999 | >0.9999 | **0.0097** |
| NBD | CTR - RHEB-p.P37L | >0.9999 | >0.9999 | >0.9999 | >0.9999 | >0.9999 | >0.9999 |
|  | *Tsc1<sup>fl/fl</sup>*-NT - *Tsc1<sup>fl/fl</sup>*-CTR | >0.9999 | 0.9718 | **0.0168** | **<0.0001** | 0.0983 | >0.9999 |
|  | *Tsc1<sup>fl/fl</sup>*-NT - *Tsc1<sup>fl/fl</sup>*-CRE | >0.9999 | >0.9999 | >0.9999 | **<0.0001** | >0.9999 | >0.9999 |
|  | *Tsc1<sup>fl/fl</sup>*-CTR - *Tsc1<sup>fl/fl</sup>*-CRE | >0.9999 | >0.9999 | **0.0014** | >0.9999 | >0.9999 | >0.9999 |
| NBC | CTR - RHEB-p.P37L | >0.9999 | >0.9999 | >0.9999 | **0.0013** | **0.0002** | **<0.0001** |
|  | *Tsc1<sup>fl/fl</sup>*-NT - *Tsc1<sup>fl/fl</sup>*-CTR | >0.9999 | >0.9999 | >0.9999 | **<0.0001** | **0.0014** | >0.9999 |
|  | *Tsc1<sup>fl/fl</sup>*-NT - *Tsc1<sup>fl/fl</sup>*-CRE | >0.9999 | >0.9999 | >0.9999 | **<0.0001** | **<0.0001** | **<0.0001** |
|  | *Tsc1<sup>fl/fl</sup>*-CTR - *Tsc1<sup>fl/fl</sup>*-CRE | >0.9999 | >0.9999 | >0.9999 | >0.9999 | >0.9999 | **<0.0001** |
| Reverberation duration | CTR - RHEB-p.P37L | 0.0714 | **0.0409** | **0.0424** | **<0.0001** | **<0.0001** | **<0.0001** |
|  | *Tsc1<sup>fl/fl</sup>*-NT - *Tsc1<sup>fl/fl</sup>*-CTR | >0.9999 | **0.0008** | >0.9999 | >0.9999 | >0.9999 | >0.9999 |
|  | *Tsc1<sup>fl/fl</sup>*-NT - *Tsc1<sup>fl/fl</sup>*-CRE | >0.9999 | >0.9999 | **<0.0001** | **0.0093** | **<0.0001** | **<0.0001** |
|  | *Tsc1<sup>fl/fl</sup>*-CTR - *Tsc1<sup>fl/fl</sup>*-CRE | >0.9999 | 0.1198 | **0.0255** | >0.9999 | **0.0044** | **<0.0001** |

MFR = Mean firing rate, %RS = percentage random spikes, NIBI = network inter-burst interval, CoV<sup>NIBI</sup> = Coefficient of variance of NIBI, NBR = Network burst rate, NBD = Network burst duration, NBC = Network burst composition.

Early in the development of cultures, network bursts occurred at a low frequency and in more irregular intervals. With more maturity of the cultures, network bursts started to occur more rhythmically (Fig 2B) (19). In RHEB-p.P37L cultures we observed shorter network interburst intervals (NIBI) at DIV07 and DIV11 compared to control cultures (Fig 2F_1_), which corresponded to the faster synchronization of the network. While in mature cultures, there was no significant difference in the NIBI between RHEB-p.P37L and their control cultures, the rhythmicity of network activity was significantly decreased, as illustrated by an increase in the coefficient of variance of the NIBI (CoV^NIBI^) from DIV14 onwards (Fig 2G_1_). In contrast to synchronicity being unaffected by expression of RHEB-WT, these cultures did fire NBs at more irregular intervals, similar to RHEB-p.P37L cultures (Fig S1D-E). In the *Tsc1*-KO cultures, the effect on network rhythmicity was less pronounced. While there was a significant reduction in NIBI in *Tsc1*-KO cultures compared to the non-transduced controls (NT) at DIV11, 14, and 18, there was no significant difference compared to control-transduced cultures (CTR) (Fig 2F_4_). Furthermore, we only detected a slight decrease in rhythmicity at DIV21 in *Tsc1*-KO cultures when the CoV^NIBI^ was increased compared to controls (Fig 2G_5_). All statistics are presented in Tables 1A-B and S1A-B.

Taken together, hyperactivity of mTORC1 resulted in increased firing rates and faster synchronization of the network. Additionally, mature cultures showed decreased rhythmicity of network burst firing, an effect that was more pronounced in the RHEB-p.P37L cultures compared to *Tsc1*-KO cultures.

### 2.2 Alterations in network burst characteristics as a result of hyperactivity of mTORC1

Next, we zoomed in on single network bursts (NBs) and analyzed their characteristics (Fig 3A). When NBs started to appear in the first to the second week, they consisted of a short sequence of high-frequency spiking activity (Fig 3B). During the second week, these bursts gained a more refined characteristic in which several reverberations of short, high-frequency spiking patterns, clustered together into a network burst (Fig 3A). These reverberations occurred in the theta frequency range (4-10Hz), which was reported to be specific to hippocampal cultures (19). We investigated differences in these patterns in the different conditions for three NB characteristics: network burst duration, network burst composition, and reverberation duration.

**Figure 3.**
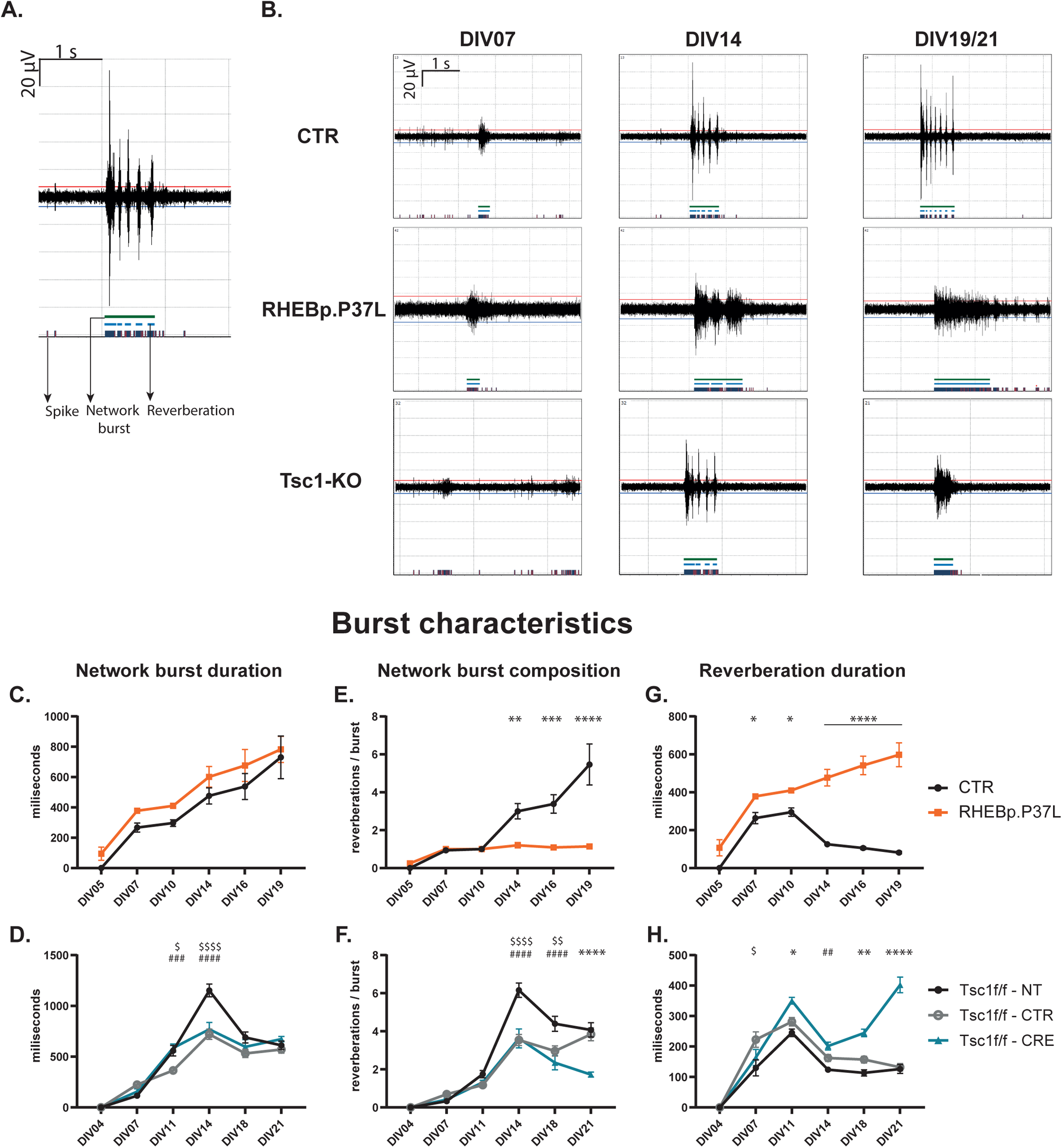
Burst characteristics are altered upon activation of mTORC1 signaling. (A) Example trace of activity recorded on a single electrode. Spikes are marked by blue-red lines at the bottom of the graph, network reverberations are marked by light blue bars (middle), and network bursts are marked by green bars (top). (B) Representative burst trains recorded on a single electrode for control, RHEB-p.P37L, and *Tsc1*-KO cultures at 3 different time points (DIV07, DIV14, and DIV19 (CTR & RHEB-p.P37L) / DIV21 (*Tsc1*-KO)). (C-H) Quantification of burst characteristics in RHEB-p.P37L and *Tsc1*-KO cultures compared to their controls: Network burst duration in milliseconds (C-D), network burst composition in reverberations/burst (E-F), and reverberation duration in milliseconds (G-H). Data is presented as mean ± SEM. Sample size (wells/replicates): CTR: n = 15 / 3, RHEB-p.P37L: n = 17 / 3. *Tsc1*^f/f^-NT: n = 42 / 4, *Tsc1*^f/f^-CTR: n = 46 / 4, *Tsc1*^f/f^-CRE: n = 51 / 4. Significant differences were determined using a Mixed-effect analysis with post hoc Bonferroni correction. *p < 0.05, **p < 0.01, ***p < 0.001, ****p < 0.0001. (D, F, H) * represent significant differences between *Tsc1*^f/f^-CRE and both controls, # represent significant differences between *Tsc1*^f/f^-CRE and either of the controls, $ represent significant differences between *Tsc1*^f/f^-NT and *Tsc1*^f/f^-CTR. All statistical comparisons are presented in Table 1A-B.

No significant changes in the duration of NBs were observed in mTORC1-hyperactive cultures (Fig 3C-D, Fig S1F). However, the induction of mTORC1 hyperactivity drastically altered the composition of the bursts. While in control cultures, mature bursts consisted of multiple reverberations, mTORC1 hyperactive cultures lost this characteristic. Instead, network bursts generally consisted of a single reverberation with a significantly longer duration (Figs 3A, E-H). In the RHEB-p.P37L cultures, this effect became apparent from DIV14 onwards (Fig 3E, G). Expression of the wildtype RHEB protein in neuronal cultures resulted in an intermediate phenotype (Fig S1G-H). While in *Tsc1*-KO cultures NBs initially consisted of several reverberations, this pattern started to change at DIV18, when *Tsc1*-KO cultures showed a lower NB composition compared to NT controls. At DIV21 this difference was more pronounced, as *Tsc1*-KO cultures showed a decreased NB composition compared to both NT and CTR-virus controls (Fig 3F, H). All statistics are presented in Tables 1A-B and S1A-B.

Overall, this data shows that in response to mTORC1 hyperactivity, burst characteristics develop differently, with the biggest alterations in mature cultures when NB composition alters and reverberation duration increases compared to controls.

### 2.3 Early rapamycin treatment of mTORC1 hyperactive cultures prevents the development of network and burst phenotypes

To investigate if the phenotypes in mTORC1-hyperactive culture recorded using the multiwell MEA system were mTORC1 hyperactivity specific, we treated the neuronal cultures with the mTORC1-inhibitor rapamycin. Rapamycin was added to the cultures early in the development of the network: in RHEB-p.P37L cultures at DIV03, and *Tsc1*-KO cultures at DIV04 (Fig 4). We found that early rapamycin treatment was able to prevent the increase in MFR and faster synchronization of the network in RHEB-p.P37L cultures (Fig 4A_1,3,5_). For the *Tsc1*-KO cultures, where we observed a more synchronous network at DIV11 and 14 in the previous phenotyping experiment, the difference was no longer observed in this experiment as both *Tsc1*-KO cultures behaved similarly to the corresponding control-treated cultures, suggesting that this parameter is not very robust (Fig 4A_4,6_). However, at DIV11 and DIV14, the network in rapamycin-treated *Tsc1*-KO cultures was less synchronous with a decreased NBR and increased %RS compared to the DMSO-treated *Tsc1*-KO cultures (Fig 4A_4,6_). The decreased NIBI observed in mTORC1 hyperactive cultures early in development was also corrected by rapamycin treatment, paralleled by the prevention of the early increase and later decrease in rhythmicity in these cultures (Fig 4B_1-4_).

**Figure 4.**
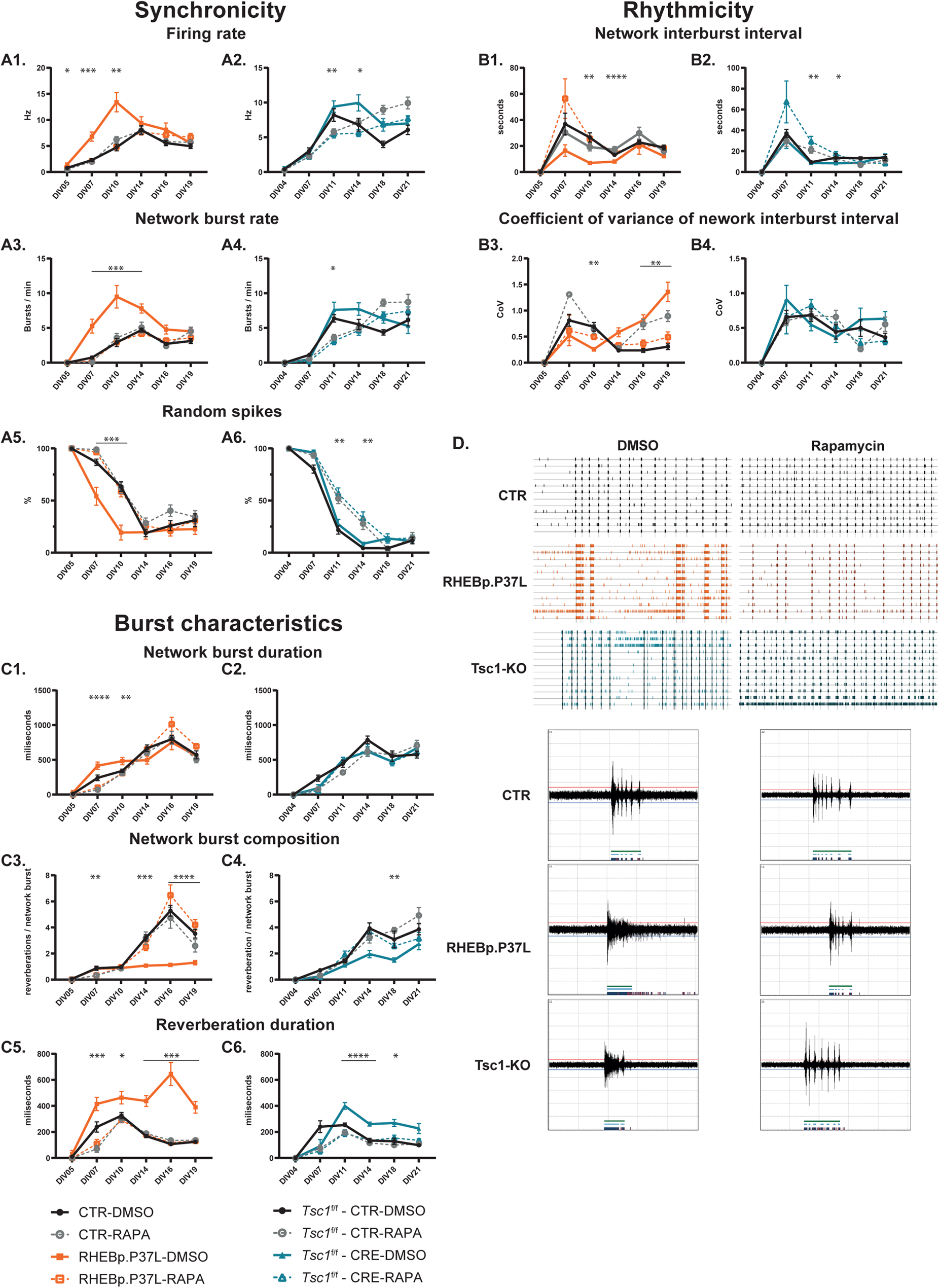
Early rapamycin treatment rescues alterations in network and burst activity in mTORC1 hyperactive cultures. Cultures were treated with rapamycin starting at DIV03 in RHEB-p.P37L cultures and DIV04 in *Tsc1*^f/f^ cultures. (A) Effect of early rapamycin treatment on network synchronicity: firing rate (A1-2), network burst rate (A3-4), and percentage random spikes (A5-6) in mTORC1 hyperactive cultures. (B) Effect of early rapamycin treatment on network rhythmicity: NIBI (B1-2), and coefficient of variance of NIBI (B3-4) in mTORC1 hyperactive cultures. (C) Effect of early rapamycin treatment on burst characteristics: network burst duration (C1-2), network burst composition (C3-4) and reverberation duration (C5-6) in mTORC1 hyperactive cultures. (D) Representative raster plots and bursts from DIV21 CTR (black), DIV19 RHEB (orange), and DIV21 *Tsc1*-KO (blue) cultures, treated early in development with DMSO (left) or rapamycin (right). Data is presented as mean ± SEM. Sample size (wells/replicates): CTR-DMSO: n = 23 / 4, CTR-RAPA: n = 22 / 4, RHEB-p.P37L-DMSO: n = 18 / 4, RHEB-p.P37L-RAPA: n = 21 / 4. *Tsc1*^f/f^-CTR-DMSO: n = 14 / 4, *Tsc1*^f/f^-CTR-RAPA: n = 19 / 4, *Tsc1*^f/f^-CRE-DMSO: n = 12 / 4, *Tsc1*^f/f^-CRE-RAPA: n = 20 / 4. Significant differences were determined using a Mixed-effect analysis with post hoc Bonferroni correction. * represent significant differences between DMSO and rapamycin treated mTORC1 hyperactive groups (RHEB-p.P37L-DMSO vs RHEB-p.P37L-RAPA and *Tsc1*^f/f^-CRE-DMSO vs *Tsc1*^f/f^-CRE-RAPA). *p < 0.05, **p < 0.01, ***p < 0.001, ****p < 0.0001. All statistical comparisons are presented in Tables 2A-C and S2A-B.

Changes in burst characteristics identified in mTORC1 hyperactive cultures could be prevented by early treatment with rapamycin. Rapamycin slightly reduced the NB duration in RHEB-p.P37L cultures compared to DMSO, but it did not affect NB duration in *Tsc1*-KO cultures (Fig 4C_1,2_). More importantly, rapamycin was able to restore the NB composition, as the rapamycin-treated mTORC1 hyperactive cultures established NBs consisting of multiple reverberations, that were of a duration similar to reverberations in control cultures (Fig 4C_3-6_). The prevention of network activity phenotypes by early rapamycin treatment is illustrated in the example traces in Fig 4D. All statistics are presented in Tables 2A-B and S2A-B, and a summary of the results is presented in Table 3.

Thus, our results show that we have identified robust phenotypes in both mTORC1 hyperactivity models. Additionally, we found that these parameters are sensitive to rapamycin treatment as early rapamycin treatment fully prevented the network and burst phenotypes we identified in mTORC1 hyperactive cultures. This confirms the mTORC1 dependency of the identified network changes.

### 2.4 Late rapamycin treatment cannot rescue all network and burst phenotypes in mTORC1 hyperactive cultures

To investigate if rapamycin is also able to rescue the identified phenotypes when treatment is started in mature cultures, in which neurons have been able to establish most of the connections in their network, we treated RHEB-p.P37L cultures at DIV10 and *Tsc1*-KO cultures at DIV14 (Fig 5). Since phenotypes related to network synchronicity occur in mTORC1 hyperactive cultures mostly before the start of the treatment, we did not find any effects of the treatment here. We did however detect a slight increase in the %RS occurring in RHEB-p.P37L after rapamycin treatment and a decrease in network burst rate at DIV14 (Fig 5A_1-6_). Additionally, the decrease in rhythmicity measured as an increase in the CoV^NIBI^, appearing from DIV14 in the RHEB-p.P37L cultures, and DIV21 in *Tsc1-*KO cultures could be rescued by late treatment with rapamycin (Fig 5B_1-4_). Interestingly, not all changes in burst characteristics could be rescued (Fig 5C). While we did find a full rescue of the reverberation duration, from DIV14 in RHEB-p.P37L cultures, and at DIV21 in *Tsc1*-KO cultures (Fig 5C_5,6_), late rapamycin treatment did not restore the NB composition. In treated mTORC1 hyperactive cultures NBs consisted of only one or few reverberations (Fig 5C_3,4_), which is also reflected by the decreased duration of NBs after late rapamycin treatment (Fig 5C_1,2_). The effect of late rapamycin treatment on network activity is illustrated in the example traces in Fig 5D. All statistics are presented in Tables 2A-C and S2A-B and a summary of the results is presented in Table 3.

**Figure 5.**
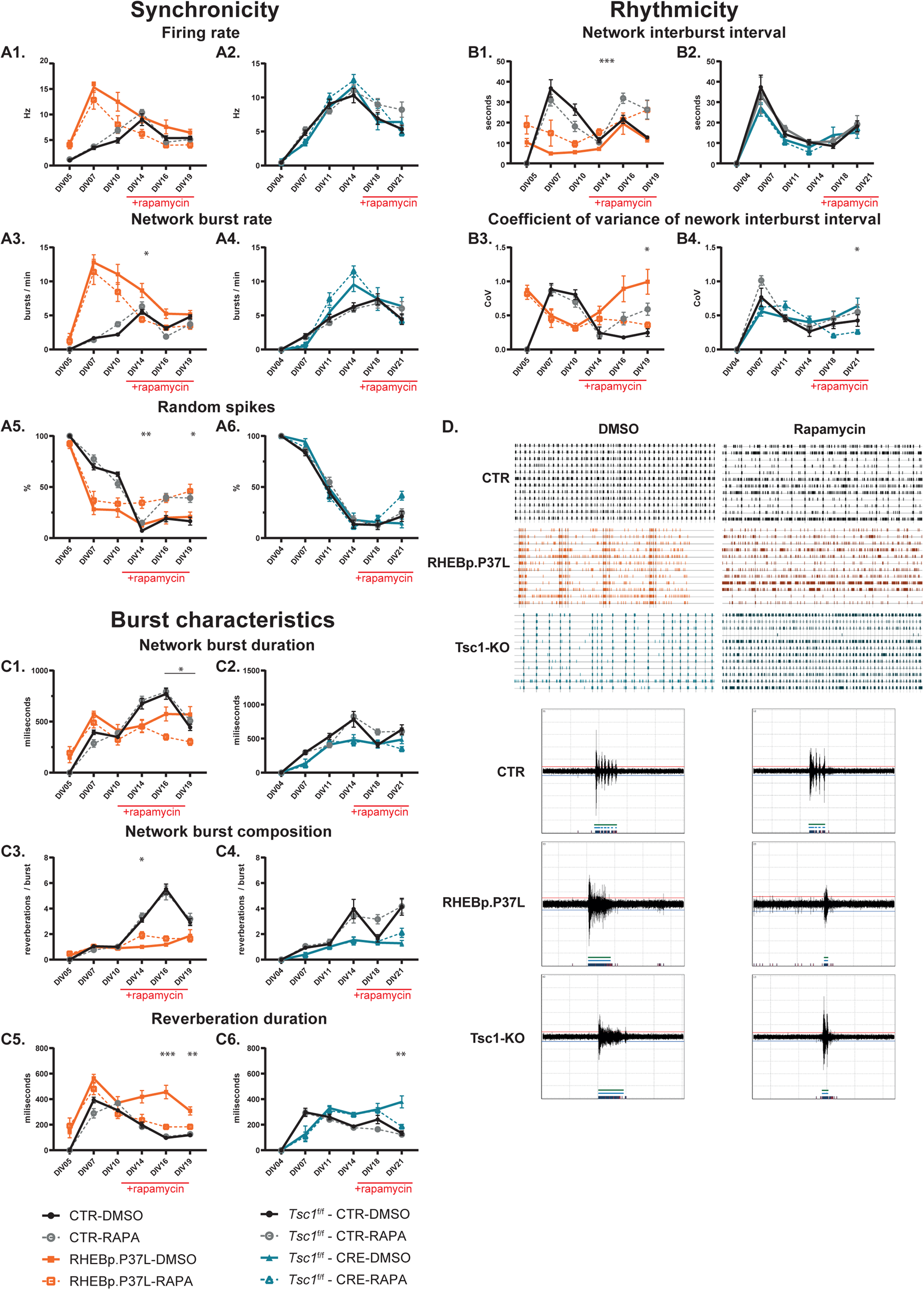
Late rapamycin treatment cannot rescue all phenotypes identified in mTORC1 hyperactive cultures. Cultures were treated with rapamycin starting at DIV10 in RHEB-p.P37L cultures and DIV14 in *Tsc1*^f/f^ cultures (A) Effect of late rapamycin treatment on network synchronicity: firing rate (A1-2), network burst rate (A3-4), and percentage random spikes (A5-6) in mTORC1 hyperactive cultures. (B) Effect of late rapamycin treatment on network rhythmicity: NIBI (B1-2) and coefficient of variance of NIBI (B3-4) in mTORC1 hyperactive cultures. (C) Effect of late rapamycin treatment on burst characteristics: network burst duration (C1-2) network burst composition (C3-4), and reverberation duration (C5-6) in mTORC1 hyperactive cultures. (D) Representative raster plots and bursts from DIV 19 CTR (black), DIV19 RHEB (orange), and DIV21 *Tsc1*-KO (blue) cultures, after late treatment with DMSO (left) or rapamycin (right). Data is presented as mean ± SEM. Sample size (wells/replicates): CTR-DMSO: n = 20 / 4, CTR-RAPA: n = 21 / 4, RHEB-p.P37L-DMSO: n = 19 / 4, RHEB-p.P37L-RAPA: n = 17 / 4. *Tsc1*^f/f^-CTR-DMSO: n = 14 / 5, *Tsc1*^f/f^-CTR-RAPA: n = 18 / 5, *Tsc1*^f/f^-CRE-DMSO: n = 15 / 5, *Tsc1*^f/f^-CRE-RAPA: n = 20 / 5. Significant differences were determined using a Mixed-effect analysis with post hoc Bonferroni correction. * represent significant differences between DMSO and rapamycin-treated mTORC1 hyperactive groups from the moment of rapamycin treatment (RHEB-p.P37L-DMSO vs RHEB-p.P37L-RAPA from DIV14 and *Tsc1*^f/f^-CRE-DMSO vs *Tsc1*^f/f^-CRE-RAPA from DIV18). *p < 0.05, **p < 0.01, ***p < 0.001, ****p < 0.0001. All statistical comparisons are presented in Table 2A-C and Supplementary table 2A-B.

**Table 2A.** Statistical analysis overview of mixed effects analysis for early and late rescue of mTORC1 hyperactivity-induced phenotypes on the MEA.

| Outcome parameter | Between effect |  | Interaction effect |  |
| --- | --- | --- | --- | --- |
|  | p-value | F | p value | F |
| **Early treatment - RHEB-p.P37L** |  |  |  |  |
| MFR | ****<0.0001 | (3, 80) = 9.950 | ****<0.0001 | (15, 388) = 7.459 |
| %RS | ****<0.0001 | (3, 80) = 11.03 | ****<0.0001 | (15, 388) = 5.374 |
| NBR | ****<0.0001 | (3, 80) = 32.75 | ****<0.0001 | (15, 388) = 5.846 |
| NIBI | ****<0.0001 | (3, 80) = 14.23 | ****<0.0001 | (15, 323) = 3.257 |
| CoV<sup>NIBI</sup> | ****<0.0001 | (3, 80) = 10.89 | ****<0.0001 | (15, 323) = 10.53 |
| NBD | *0.0475 | (3, 80) = 2.760 | ****<0.0001 | (15, 390) = 3.797 |
| NBC | ****<0.0001 | (3, 80) = 19.33 | ****<0.0001 | (15, 389) = 8.832 |
| Reverberation duration | ****<0.0001 | (3, 80) = 104.0 | ****<0.0001 | (15, 388) = 8.865 |
| **Late treatment - RHEB-p.P37L** |  |  |  |  |
| MFR | ****<0.0001 | (3, 73) = 17.17 | ****<0.0001 | (15, 349) = 10.98 |
| %RS | ****<0.0001 | (3, 422) = 22.44 | ****<0.0001 | (15, 422) = 9.640 |
| NBR | ****<0.0001 | (3, 73) = 53.53 | ****<0.0001 | (15, 349) = 13.11 |
| NIBI | ****<0.0001 | (3, 73) = 10.35 | ****<0.0001 | (15, 313) = 7.591 |
| CoV<sup>NIBI</sup> | ***0.0001 | (3, 73) = 8.015 | ****<0.0001 | (15, 313) = 10.59 |
| NBD | **0.0074 | (3, 73) = 4.311 | ****<0.0001 | (15, 349) = 9.803 |
| NBC | ****<0.0001 | (3, 73) = 22.46 | ****<0.0001 | (15, 349) = 18.42 |
| Reverberation duration | ****<0.0001 | (3, 73) = 49.72 | ****<0.0001 | (15, 349) = 4.874 |
| **Early treatment - *Tsc1*-KO** |  |  |  |  |
| MFR | *<0.0494 | (3, 61) = 2.766 | ****<0.0001 | (15, 293) = 8.183 |
| %RS | ****<0.0001 | (3, 61) = 10.16 | ****<0.0001 | (15, 293) = 5.585 |
| NBR | 0.1260 | (3, 61) = 1.982 | ****<0.0001 | (15, 293) = 6.285 |
| NIBI | ****<0.001 | (3, 61) = 10.58 | ****<0.0001 | (15, 250) = 5.825 |
| CoV<sup>NIBI</sup> | 0.5042 | (3, 311) = 0.7829 | ***0.0002 | (15, 311) = 2.948 |
| NBD | 0.3402 | (3, 61) = 1.140 | **0.0081 | (15, 293) = 2.150 |
| NBC | ***0.0002 | (3, 61) = 7.894 | ***0.0009 | (15, 293) = 2.653 |
| Reverberation duration | ****<0.0001 | (3, 61) = 13.63 | ****<0.0001 | (15, 293) = 7.763 |

| Late treatment - *Tsc1*-KO |  |  |  |  |
| --- | --- | --- | --- | --- |
| MFR | 0.3595 | (3, 63) = 1.091 | ****<0.0001 | (15, 304) = 3.586 |
| %RS | 0.5709 | (3, 63) = 0.6745 | ****<0.0001 | (15, 304) = 3.197 |
| NBR | *0.0127 | (3, 63) = 3.902 | ****<0.0001 | (15, 304) = 4.715 |
| NIBI | 0.0886 | (3, 63) = 2.274 | 0.7840 | (15, 272) = 0.7000 |
| CoV<sup>NIBI</sup> | *0.0324 | (3, 335) = 2.961 | ****<0.0001 | (15, 335) = 3.235 |
| NBD | ****<0.0001 | (3, 63) = 15.35 | ****<0.0001 | (15, 304) = 3.577 |
| NBC | ****<0.0001 | (3, 367) = 27.06 | ****<0.0001 | (15, 367) = 4.519 |
| Reverberation duration | ***0.0006 | (3, 63) = 6.546 | ****<0.0001 | (15, 304) = 10.40 |

MFR = Mean firing rate, %RS = percentage random spikes, NIBI = network inter-burst interval, CoV<sup>NIBI</sup> = Coefficient of variance of NIBI, NBR = Network burst rate, NBD = Network burst duration, NBC = Network burst composition.

**Table 2B.** Post-hoc Bonferroni corrected multiple comparisons for early and late rescue of RHEB-p.P37L induced mTORC1 hyperactivity phenotypes on the MEA.

| Parameter | Comparison | p-value |  |  |  |  |  |
| --- | --- | --- | --- | --- | --- | --- | --- |
|  |  | DIV04/05 | DIV07 | DIV10/11 | DIV14 | DIV16/18 | DIV19/21 |
| Early treatment – RHEB-p.P37L |  |  |  |  |  |  |  |
| MFR | CTR-DMSO – CTR-RAPA | 0.1135 | >0.9999 | 0.5783 | >0.9999 | >0.9999 | >0.9999 |
|  | CTR-DMSO – RHEB-p.P37L-RAPA | >0.9999 | >0.9999 | >0.9999 | >0.9999 | 0.4791 | 0.3071 |
|  | RHEB-p.P37L -DMSO – RHEB-p.P37L-RAPA | **0.0144** | **0.0003** | **0.0015** | >0.9999 | >0.9999 | >0.9999 |
| %RS | CTR-DMSO – CTR-RAPA | >0.9999 | **0.0038** | >0.9999 | 0.6884 | 0.3320 | >0.9999 |
|  | CTR-DMSO – RHEB-p.P37L-RAPA | >0.9999 | **0.0485** | >0.9999 | >0.9999 | >0.9999 | >0.9999 |
|  | RHEB-p.P37L -DMSO – RHEB-p.P37L- RAPA | >0.9999 | **0.0007** | **0.0005** | >0.9999 | >0.9999 | >0.9999 |
| NBR | CTR-DMSO – CTR-RAPA | >0.9999 | **0.0126** | >0.9999 | >0.9999 | >0.9999 | 0.2916 |
|  | CTR-DMSO – RHEB-p.P37L-RAPA | >0.9999 | 0.2831 | >0.9999 | >0.9999 | 0.9006 | >0.9999 |
|  | RHEB-p.P37L -DMSO – RHEB-p.P37L- RAPA | >0.9999 | **0.0006** | **0.0059** | **0.0004** | 0.2129 | >0.9999 |
| NIBI | CTR-DMSO – CTR-RAPA |  |  | 0.6844 | 0.7776 | 0.9411 | >0.9999 |
|  | CTR-DMSO – RHEB-p.P37L-RAPA |  | >0.9999 | >0.9999 | >0.9999 | >0.9999 | >0.9999 |
|  | RHEB-p.P37L -DMSO – RHEB-p.P37L- RAPA |  | 0.4361 | **0.0029** | **<0.0001** | >0.9999 | 0.1100 |
| CoV<sup>NIBI</sup> | CTR-DMSO – CTR-RAPA |  |  | >0.9999 | >0.9999 | **0.0004** | **0.0001** |
|  | CTR-DMSO – RHEB-p.P37L-RAPA |  | >0.9999 | 0.4294 | 0.2804 | 0.7157 | 0.7587 |
|  | RHEB-p.P37L -DMSO – RHEB-p.P37L- RAPA |  | >0.9999 | **0.0062** | 0.0944 | **0.0074** | **0.0022** |
| NBD | CTR-DMSO – CTR-RAPA | >0.9999 | **0.0060** | >0.9999 | >0.9999 | >0.9999 | >0.9999 |
|  | CTR-DMSO – RHEB-p.P37L-RAPA | >0.9999 | 0.0599 | >0.9999 | >0.9999 | 0.4089 | 0.5232 |
|  | RHEB-p.P37L -DMSO – RHEB-p.P37L- RAPA | >0.9999 | **<0.0001** | **0.0268** | 0.1351 | 0.4843 | 0.6396 |
| NBC | CTR-DMSO – CTR-RAPA | >0.9999 | 0.0502 | >0.9999 | >0.9999 | >0.9999 | 0.7241 |
|  | CTR-DMSO – RHEB-p.P37L-RAPA | >0.9999 | **0.0451** | >0.9999 | 0.5728 | >0.9999 | >0.9999 |
|  | RHEB-p.P37L -DMSO – RHEB-p.P37L- RAPA | >0.9999 | **0.0056** | >0.9999 | **0.0008** | **<0.0001** | **<0.0001** |
| Reverberation duration | CTR-DMSO – CTR-RAPA | >0.9999 | **0.0065** | >0.9999 | >0.9999 | 0.0504 | >0.9999 |
|  | CTR-DMSO – RHEB-p.P37L-RAPA | >0.9999 | 0.1060 | >0.9999 | >0.9999 | >0.9999 | >0.9999 |
|  | RHEB-p.P37L -DMSO – RHEB-p.P37L- RAPA | >0.9999 | **0.0002** | **0.0174** | **0.0001** | **0.0002** | **0.0002** |
|  |  | DIV04/05 | DIV07 | DIV10/11 | DIV14 | DIV16/18 | DIV19/21 |
| Late treatment – RHEB-p.P37L |  |  |  |  |  |  |  |
| MFR | CTR-DMSO – CTR-RAPA | >0.9999 | >0.9999 | 0.0775 | >0.9999 | 0.7988 | >0.9999 |
|  | CTR-DMSO – RHEB-p.P37L-RAPA | **0.0061** | **0.0005** | 0.6086 | 0.3685 | 0.2639 | 0.4743 |
|  | RHEB-p.P37L -DMSO – RHEB-p.P37L- RAPA | >0.9999 | >0.9999 | 0.5153 | 0.2026 | 0.1343 | 0.0892 |
| %RS | CTR-DMSO – CTR-RAPA |  | 0.7149 | 0.2274 | **0.0113** | **0.0008** | **0.0021** |
|  | CTR-DMSO – RHEB-p.P37L-RAPA | 0.1733 | **0.0137** | **0.0006** | **0.0007** | **0.0019** | **0.0056** |
|  | RHEB-p.P37L -DMSO – RHEB-p.P37L- RAPA | >0.9999 | >0.9999 | >0.9999 | **0.0094** | 0.0763 | **0.0300** |
| NBR | CTR-DMSO – CTR-RAPA |  | >0.9999 | **0.0036** | >0.9999 | **0.0019** | 0.7503 |
|  | CTR-DMSO – RHEB-p.P37L-RAPA | 0.2491 | **0.0005** | **0.0021** | 0.3313 | >0.9999 | 0.2709 |
|  | RHEB-p.P37L -DMSO – RHEB-p.P37L- RAPA | >0.9999 | >0.9999 | >0.9999 | **0.0178** | 0.0836 | 0.2603 |
| NIBI | CTR-DMSO – CTR-RAPA |  | >0.9999 | 0.1946 | >0.9999 | **0.0158** | 0.0516 |
|  | CTR-DMSO – RHEB-p.P37L-RAPA | 0.0780 | **0.0483** | **<0.0001** | 0.4041 | >0.9999 | 0.0650 |
|  | RHEB-p.P37L -DMSO – RHEB-p.P37L- RAPA | 0.7954 | 0.8313 | 0.1853 | **0.0009** | >0.9999 | 0.0504 |
| **CoV<sup>NIBI</sup>** | CTR-DMSO – CTR-RAPA |  | >0.9999 | >0.9999 | >0.9999 | 0.0521 | **0.0153** |
|  | CTR-DMSO – RHEB-p.P37L-RAPA | **0.0031** | 0.0559 | **<0.0001** | 0.6291 | **<0.0001** | 0.7903 |
|  | RHEB-p.P37L -DMSO – RHEB-p.P37L- RAPA | >0.9999 | >0.9999 | >0.9999 | >0.9999 | 0.1556 | **0.0177** |
| **NBD** | CTR-DMSO – CTR-RAPA |  | 0.0958 | 0.3216 | >0.9999 | >0.9999 | >0.9999 |
|  | CTR-DMSO – RHEB-p.P37L-RAPA | **0.0406** | 0.6392 | >0.9999 | **0.0494** | **<0.0001** | **0.0225** |
|  | RHEB-p.P37L -DMSO – RHEB-p.P37L- RAPA | >0.9999 | >0.9999 | >0.9999 | >0.9999 | **0.0370** | **0.0234** |
| **NBC** | CTR-DMSO – CTR-RAPA |  | 0.0712 |  | >0.9999 | >0.9999 | >0.9999 |
|  | CTR-DMSO – RHEB-p.P37L-RAPA | **0.0406** | >0.9999 | >0.9999 | **0.0065** | **<0.0001** | **0.0071** |
|  | RHEB-p.P37L -DMSO – RHEB-p.P37L- RAPA | >0.9999 |  | >0.9999 | **0.0287** | 0.4187 | >0.9999 |
| **Reverberation duration** | CTR-DMSO – CTR-RAPA |  | 0.1275 | **0.0243** | >0.9999 | >0.9999 | >0.9999 |
|  | CTR-DMSO – RHEB-p.P37L-RAPA | **0.0423** | 0.5906 | >0.9999 | >0.9999 | **<0.0001** | **0.0020** |
|  | RHEB-p.P37L -DMSO – RHEB-p.P37L- RAPA | >0.9999 | 0.8538 | 0.9020 | 0.0695 | **0.0005** | **0.0092** |

MFR = Mean firing rate, %RS = percentage random spikes, NIBI = network inter-burst interval, CoV<sup>NIBI</sup> = Coefficient of variance of NIBI, NBR = Network burst rate, NBD = Network burst duration, NBC = Network burst composition.

**Table 2C.**
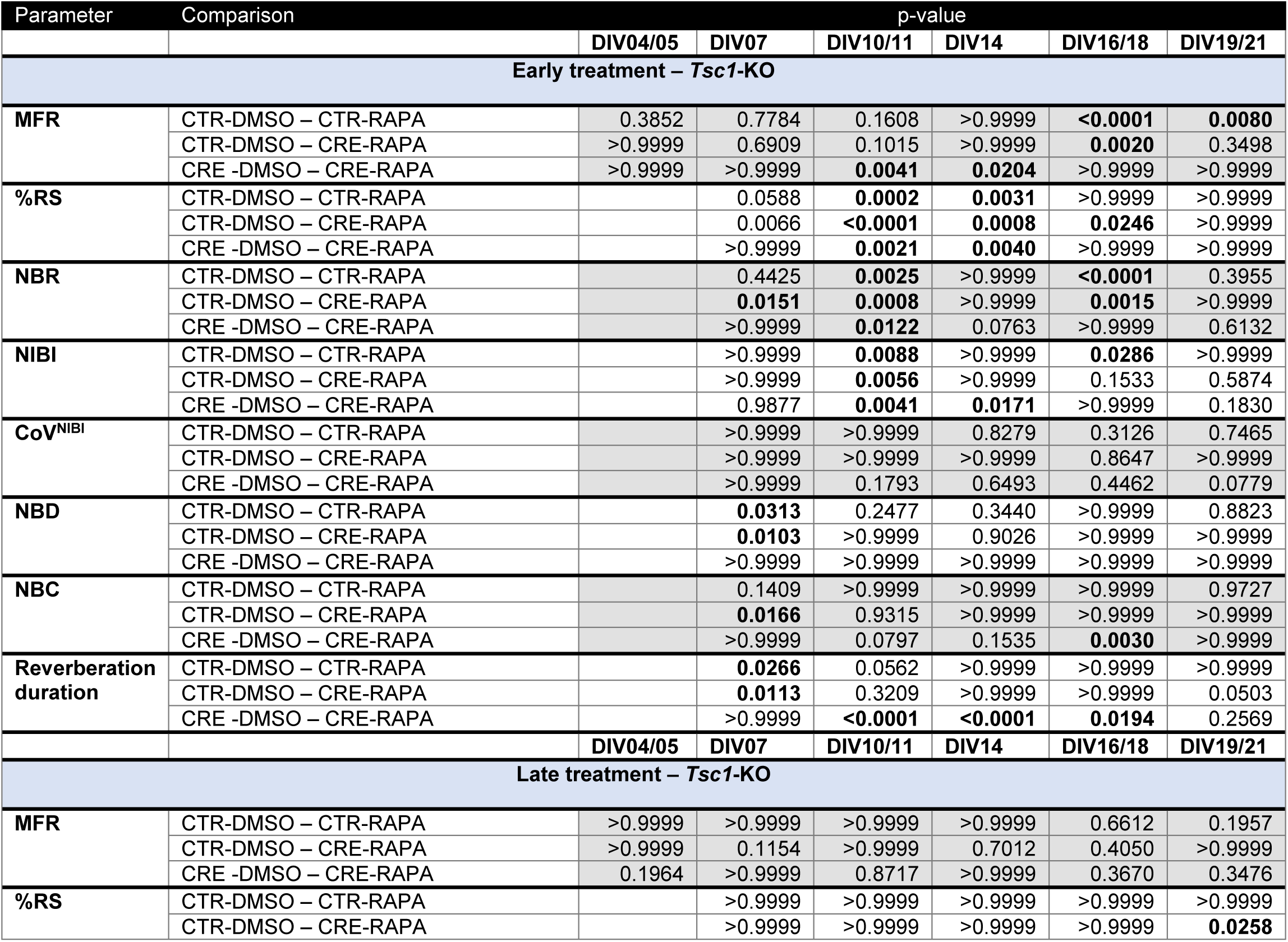
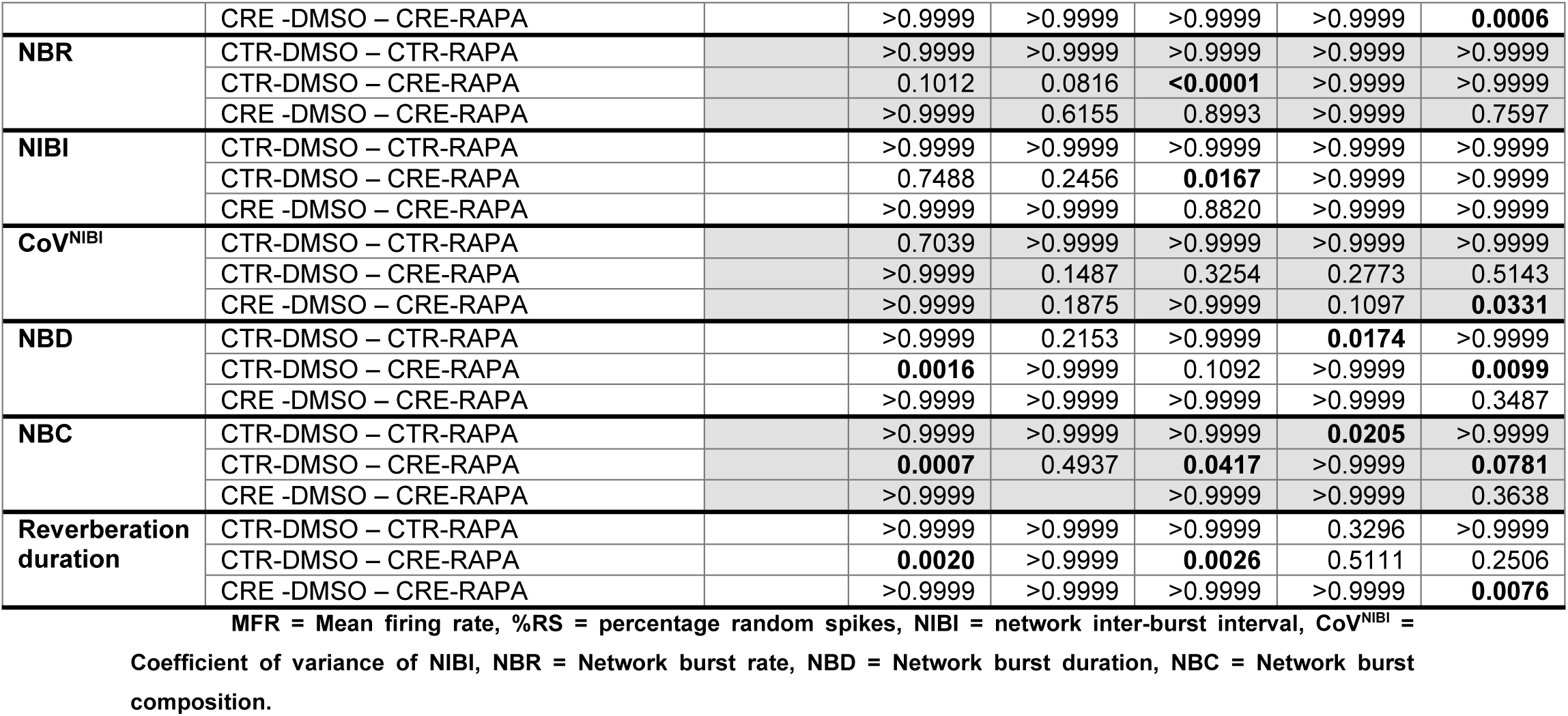
Post-hoc Bonferroni corrected multiple comparisons for early and late rescue of *Tsc1*-KO induced mTORC1 hyperactivity phenotypes on the MEA. MFR.

**Table 3.** Overview of phenotypes and the effect of treatments on each outcome parameter.

|  | Phenotype |  | Prevention / Rescue of phenotype |  |  |  |  |  |
| --- | --- | --- | --- | --- | --- | --- | --- | --- |
|  |  |  | Early rapamycin |  | Late rapamycin |  | Acute vigabatrin | Vigabatrin wash-out |
|  | RHEB-p.P37L | Tsc1-KO | RHEB-p.P37L | Tsc1-KO | RHEB-p.P37L | Tsc1-KO | RHEB-p.P37L | RHEB-p.P37L |
| **MFR** | Yes | Yes | Yes | Yes | - | - | Yes | No |
| **%RS** | Yes | Yes | Yes | - | - | - | No | No |
| **NBR** | Yes | Yes | Yes | Yes | Yes | No | Yes | No |
| **NIBI** | Yes | Yes | Yes | Yes | Yes | Yes | Yes | No |
| **CoV<sup>NIBI</sup>** | Yes | Yes | Yes | No | No | No | Yes | No |
| **NBD** | No | No | - | - | - | - | - | - |
| **NBC** | Yes | Yes | Yes | Yes | No | No | Yes | Yes |
| **Reverberation duration** | Yes | Yes | Yes | Yes | Yes | Yes | Yes | Yes |

Yes (orange/blue) = phenotype identified, Yes (green) = prevented/rescued phenotype, No = not prevented/rescued,
- = no phenotype to rescue.

Thus, although early rapamycin was able to prevent all network and burst phenotypes in mTORC1 hyperactive cultures, downregulation of mTORC1 activity in mature cultures could only rescue part of the phenotypes. Mature cultures organized their activity rhythmically upon late rapamycin treatment similar to control cultures. However, the treated cultures fired shorter bursts consisting of single reverberations instead of clustering together a sequence of multiple reverberations.

### 2.5 Treatment of RHEB-p.P37L cultures with the AED vigabatrin reduces network activity to levels similar to control activity shortly after the addition of the drug

Next, we investigated if the identified phenotypes can be used as a model for epilepsy by testing their sensitivity to the AED vigabatrin, an AED that is often prescribed to mTORopathy patients (10,30). To this extent, we used the RHEB-p.P37L cultures as in this model, the network-and burst phenotypes were most pronounced. At DIV11, when multiple mTORC1 hyperactivity-related phenotypes could be identified, we treated the cultures with vigabatrin (Fig 6A). Within one hour after vigabatrin treatment, the MFR and NBR of RHEB-p.P37L cultures were reduced to levels similar to vehicle-treated control cultures, which persisted until 5 hours after treatment (Fig 6B-C). However, in both the RHEB-p.P37L and control cultures treated with vigabatrin networks became more synchronous, as the %RS significantly decreased compared to vehicle-treated cultures (Fig 6D). Additionally, the network activity became more rhythmic in both vigabatrin-treated RHEB-p.P37L and control cultures (Fig 6E-F). Vigabatrin also affected the network burst characteristics. Reverberation duration decreased to levels similar to control cultures (Fig 6I). However, NBs already developed a reverberation pattern that is characteristic of more mature cultures, which contributes to the increase in NB duration in the RHEB-p.P37L cultures treated with vigabatrin (Fig 6G-H). A summary of the results is presented in Table 3. All statistics are presented in Tables 4A-B and S3.

**Figure 6.**
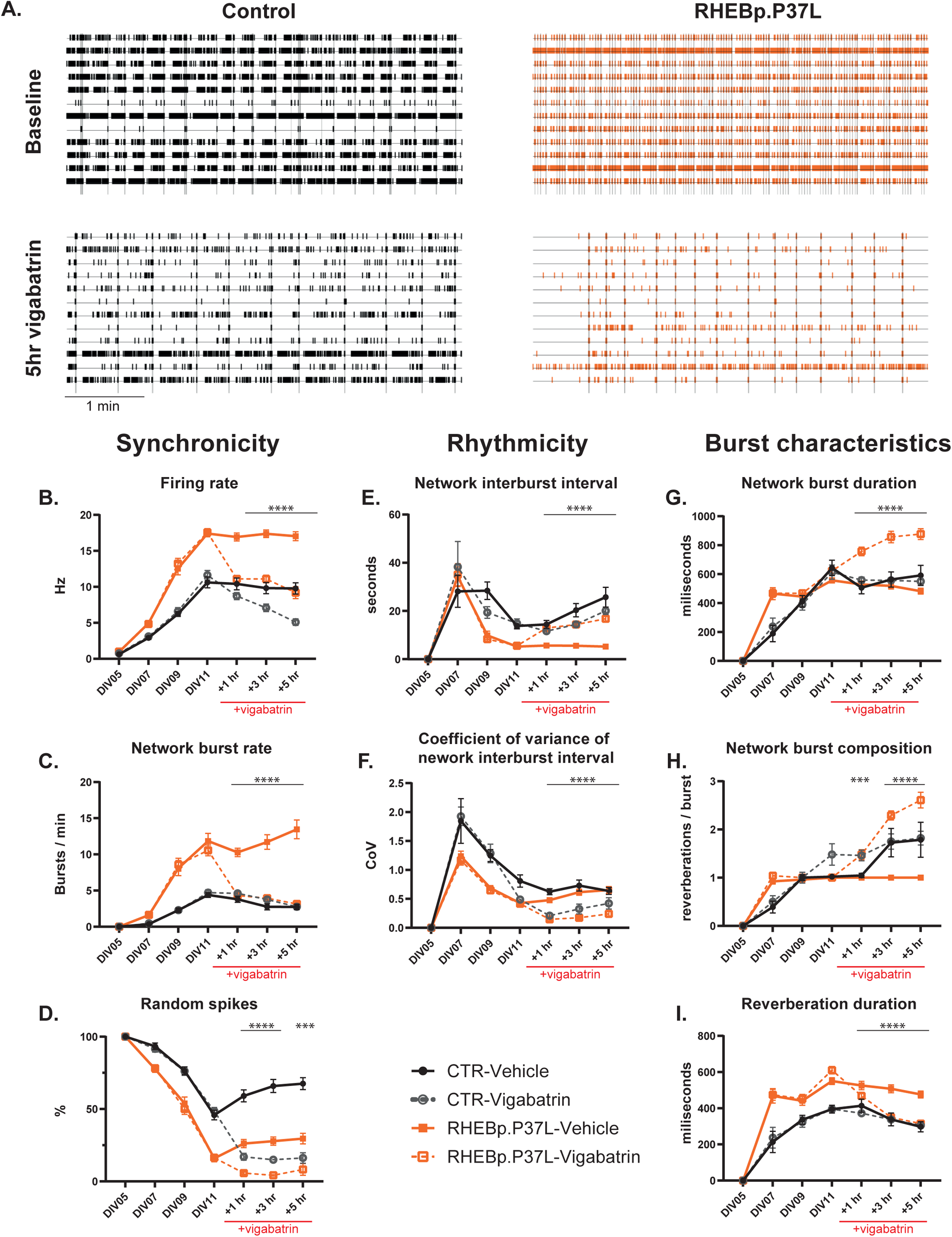
Short-term effects of vigabatrin treatment: vigabatrin treatment decreases network activity in cultures expressing RHEB-p.P37L. Cultures were treated with Vigabatrin after a baseline recording at DIV11 and the short-term effects of vigabatrin were assessed 1, 3, and 5 hours after adding vigabatrin to the cultures. (A) Representative raster plots from control-and RHEB-p.P37L-expressing cultures at DIV11 baseline (top), and 5 hours post vigabatrin treatment (bottom). (B-D) Effect of vigabatrin treatment on network synchronicity: firing rate (B), network burst rate (C), and percentage random spikes (D) in RHEB-p.P37L-expressing cultures. (E-F) Effect of vigabatrin treatment on network rhythmicity: NIBI (E) and coefficient of variance of NIBI (F) in RHEB-p.P37L-expressing cultures. (G-I) Effect of vigabatrin treatment on burst characteristics: network burst duration (G) network burst composition (H), and reverberation duration (I). Data is presented as mean ± SEM. Sample size (wells/replicates): CTR-VH: n = 24 / 2, CTR-VGB: n = 26 / 2, RHEB-p.P37L-VH: n = 26 / 2, RHEB-p.P37L-VGB: n = 24 / 2. Significant differences were determined using a Mixed-effect analysis with post hoc Bonferroni correction. * represent significant differences between vehicle (VH) and vigabatrin (VGB)-treated RHEB-p.P37L-expressing cultures. *p < 0.05, **p < 0.01, ***p < 0.001, ****p < 0.0001. All statistical comparisons are presented in Table 4A-B and S3.

**Table 4A.**
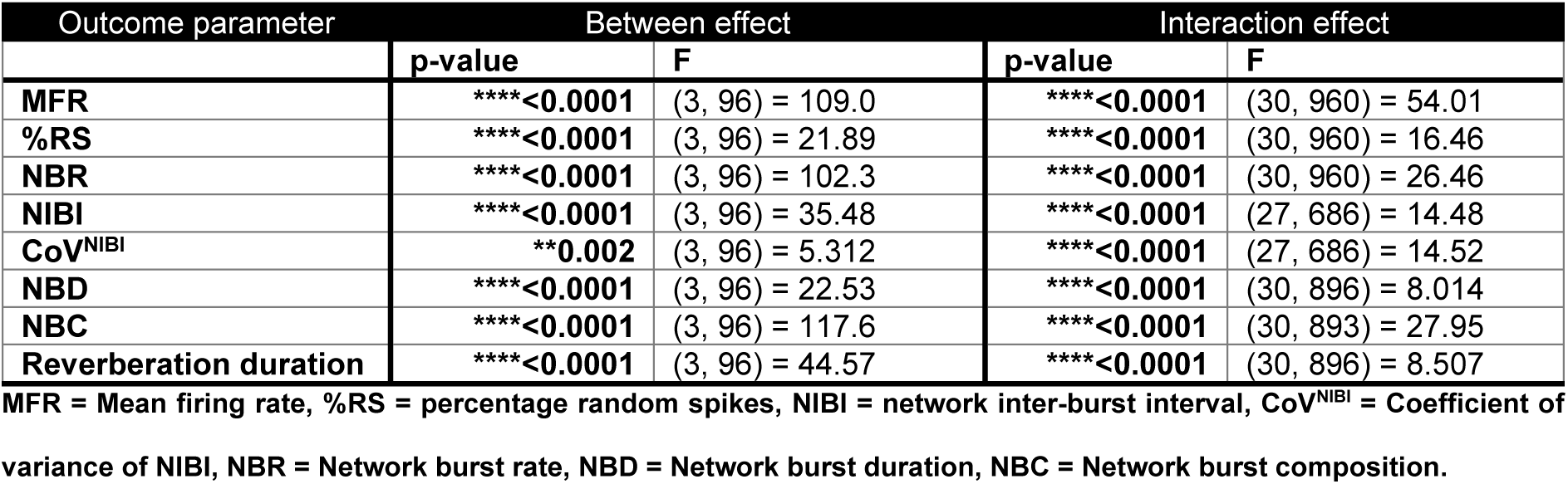
Statistical analysis overview of mixed effects analysis for vigabatrin treatment of mTORC1 hyperactivity-induced phenotypes on the MEA.

**Table 4B.** Post-hoc Bonferroni corrected multiple comparisons for vigabatrin treatment in RHEB-p.P37L-induced mTORC1 hyperactivity phenotypes on the MEA.

| Parameter | Comparison | p-value |  |  |  |  |  |  |  |  |  |  |
| --- | --- | --- | --- | --- | --- | --- | --- | --- | --- | --- | --- | --- |
|  |  | DIV05 | DIV07 | DIV09 | DIV11 | DIV11 + 1hr | DIV11 + 3hr | DIV11 + 5hr | DIV12 | DIV14 | DIV16 | DIV19 |
| **MFR** | CTR-VH – CTR-VGB | >0.9999 | >0.9999 | >0.9999 | >0.9999 | 0.4992 | **0.0344** | **<0.0001** | **<0.0001** | **<0.0001** | **<0.0001** | **<0.0001** |
|  | CTR-VH – RHEB-p.P37L-VGB | **0.0119** | **0.0004** | **<0.0001** | **<0.0001** | >0.9999 | >0.9999 | >0.9999 | **<0.0001** | **<0.0001** | **<0.0001** | **<0.0001** |
|  | RHEB-p.P37L-VH – RHEB-p.P37L-VGB | >0.9999 | >0.9999 | >0.9999 | >0.9999 | **<0.0001** | **<0.0001** | **<0.0001** | **<0.0001** | **<0.0001** | **<0.0001** | **<0.0001** |
| **%RS** | CTR-VH – CTR-VGB |  | >0.9999 | >0.9999 | >0.9999 | **<0.0001** | **<0.0001** | **<0.0001** | 0.1557 | 0.0654 | >0.9999 | 0.801 |
|  | CTR-VH – RHEB-p.P37L-VGB |  | **0.0001** | **<0.0001** | **<0.0001** | **<0.0001** | **<0.0001** | **<0.0001** | 0.071 | 0.8152 | >0.9999 | **0.0075** |
|  | RHEB-p.P37L-VH – RHEB-p.P37L-VGB |  | >0.9999 | >0.9999 | >0.9999 | **<0.0001** | **<0.0001** | **0.0019** | 0.0576 | **0.0175** | 0.4313 | **0.0073** |
| **NBR** | CTR-VH – CTR-VGB |  | >0.9999 | >0.9999 | >0.9999 | 0.6902 | 0.3923 | >0.9999 | **<0.0001** | **<0.0001** | **<0.0001** | **<0.0001** |
|  | CTR-VH – RHEB-p.P37L-VGB |  | **0.0002** | **<0.0001** | **<0.0001** | >0.9999 | 0.071 | >0.9999 | **0.0022** | **<0.0001** | **<0.0001** | **<0.0001** |
|  | RHEB-p.P37L-VH – RHEB-p.P37L-VGB |  | >0.9999 | >0.9999 | >0.9999 | **<0.0001** | **<0.0001** | **<0.0001** | **<0.0001** | **<0.0001** | **<0.0001** | **<0.0001** |
| **NIBI** | CTR-VH – CTR-VGB |  | >0.9999 | 0.2743 | >0.9999 | 0.8292 | 0.338 | >0.9999 |  | 0.1535 | **0.0266** | **0.0014** |
|  | CTR-VH – RHEB-p.P37L-VGB |  | >0.9999 | **0.0001** | **<0.0001** | >0.9999 | 0.2807 | 0.28 | 0.1021 | **<0.0001** | **0.0023** | 0.3461 |
|  | RHEB-p.P37L-VH – RHEB-p.P37L-VGB |  | >0.9999 | >0.9999 | >0.9999 | **<0.0001** | **<0.0001** | **<0.0001** | **0.0016** | **<0.0001** | **0.0007** | 0.4029 |
| **CoV<sup>NIBI</sup>** | CTR-VH – CTR-VGB |  | >0.9999 | >0.9999 | **0.0464** | **<0.0001** | **0.0165** | 0.4085 |  | 0.5894 | **<0.0001** | **0.0006** |
|  | CTR-VH – RHEB-p.P37L-VGB |  | 0.8476 | **<0.0001** | **0.0133** | **<0.0001** | **<0.0001** | **<0.0001** | **0.0058** | **0.0004** | **<0.0001** | 0.151 |
|  | RHEB-p.P37L-VH – RHEB-p.P37L-VGB |  | >0.9999 | >0.9999 | >0.9999 | **<0.0001** | **<0.0001** | **<0.0001** | 0.6837 | **0.0004** | **0.0028** | >0.9999 |
| **NBD** | CTR-VH – CTR-VGB |  | >0.9999 | >0.9999 | >0.9999 | >0.9999 | >0.9999 | >0.9999 | 0.7545 | >0.9999 | **0.0218** | **<0.0001** |
|  | CTR-VH – RHEB-p.P37L-VGB |  | 0.0006 | >0.9999 | >0.9999 | **0.0001** | **0.0004** | **0.0058** | **<0.0001** | 0.2006 | **0.0017** | 0.0659 |
|  | RHEB-p.P37L-VH – RHEB-p.P37L-VGB |  | >0.9999 | >0.9999 | 0.1663 | **<0.0001** | **<0.0001** | **<0.0001** | **<0.0001** | **<0.0001** | 0.5204 | >0.9999 |
| **NBC** | CTR-VH – CTR-VGB |  | >0.9999 | >0.9999 | 0.301 | **0.0086** | >0.9999 | >0.9999 | **0.0002** | **0.0014** | **<0.0001** | **<0.0001** |
|  | CTR-VH – RHEB-p.P37L-VGB |  | **0.0003** | >0.9999 | >0.9999 | **0.0029** | 0.4755 | 0.2878 | **0.0003** | **<0.0001** | **<0.0001** | 0.2461 |
|  | RHEB-p.P37L-VH – RHEB-p.P37L-VGB |  | >0.9999 | >0.9999 |  | **0.0006** | **<0.0001** | **<0.0001** | **0.0002** | **<0.0001** | **<0.0001** | 0.0847 |
| **Reverberation duration** | CTR-VH – CTR-VGB |  | >0.9999 | >0.9999 | >0.9999 | >0.9999 | >0.9999 | >0.9999 | >0.9999 | **<0.0001** | **<0.0001** | **<0.0001** |
|  | CTR-VH – RHEB-p.P37L-VGB |  | **0.0022** | **0.0102** | **<0.0001** | >0.9999 | >0.9999 | >0.9999 | **0.0017** | **<0.0001** | 0.052 | >0.9999 |
|  | RHEB-p.P37L-VH – RHEB-p.P37L-VGB |  | >0.9999 | >0.9999 | 0.2042 | 0.2797 | **<0.0001** | **<0.0001** | **<0.0001** | **<0.0001** | **<0.0001** | 0.8296 |

MFR = Mean firing rate, %RS = percentage random spikes, NIBI = network inter-burst interval, CoV<sup>NIBI</sup> = Coefficient of variance of NIBI, NBR = Network burst rate, NBD = Network burst
duration, NBC = Network burst composition.

### 2.6 Vigabatrin treatment of RHEB-p.P37L cultures reduces network activity and introduces the firing of multiple reverberations in a network burst

24 hours after the addition of vigabatrin to the culture, network activity was strongly suppressed in all cultures treated with vigabatrin, as illustrated by a dramatic decrease in both the MFR and NBR (Fig 7B-C). While most wells fired only a few network bursts within the 5-minute recording, in some wells no network bursts were observed at all. Since the effect of vigabatrin was so strong, we performed a wash-out experiment by gradually decreasing the vigabatrin concentration after each recording, starting with the medium refreshment on DIV12. Interestingly, in vigabatrin-treated wells that did show bursting activity, NBs consisted of significantly more reverberations than the vehicle-treated cultures and reverberation duration significantly decreased compared to the non-treated RHEB-p.P37L cultures (Fig 7H-I). A summary of the results is presented in Table 3. All statistics are presented in Tables 4A-B and S3.

**Figure 7.**
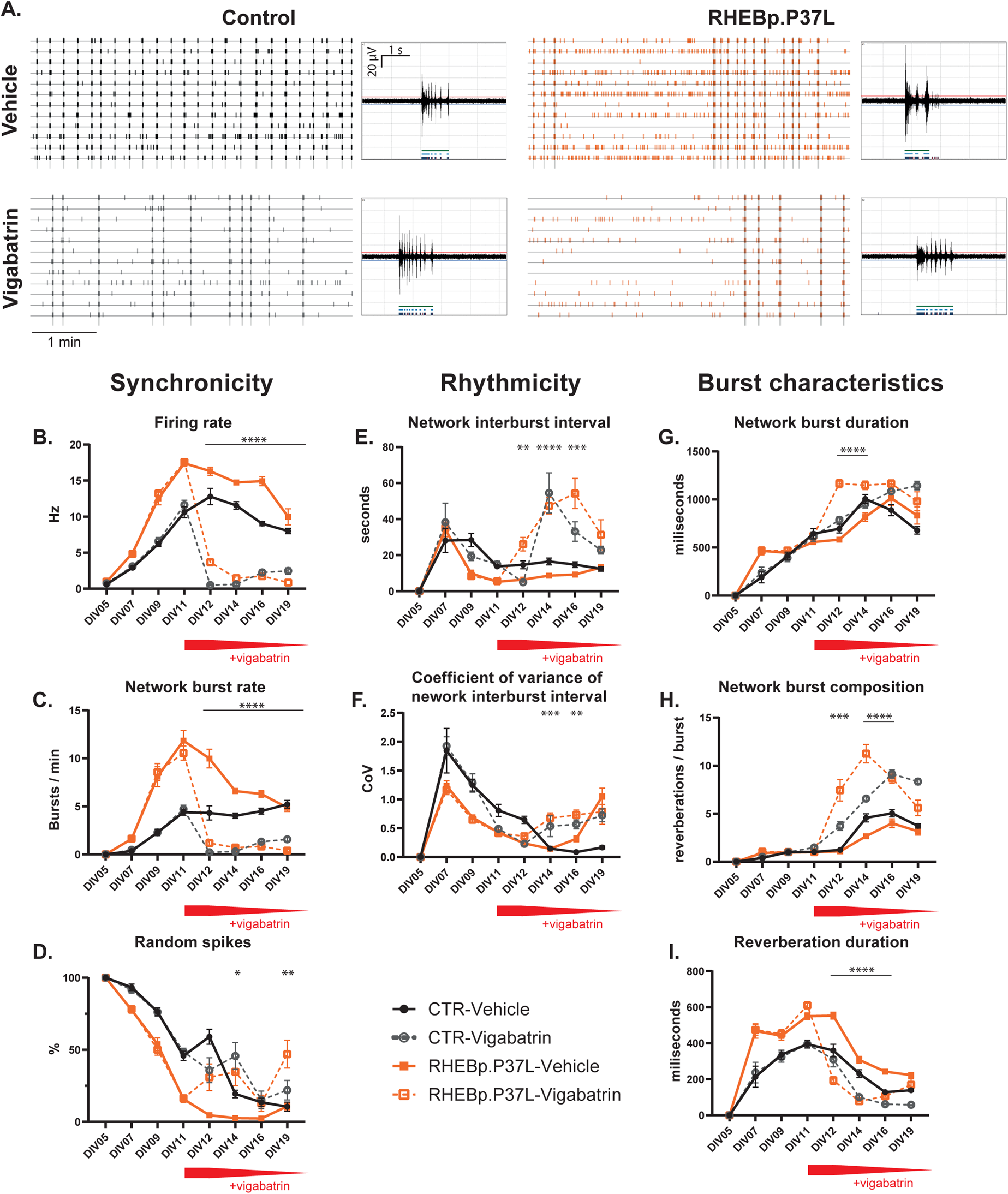
Persisting effects of vigabatrin treatment with gradual wash-out of the drug: vigabatrin treatment decreases network activity and alters burst characteristics in cultures expressing RHEB-p.P37L. Cultures were treated with vigabatrin after a baseline recording at DIV11. The persisting effects of vigabatrin were assessed over 1 week after adding vigabatrin to the cultures, while gradually washing out the drug. Representative raster plots and bursts from control-and RHEB-p.P37L-expressing cultures at DIV19 treated with vehicle (top) or vigabatrin (bottom) at DIV11. (B-D) Effect of vigabatrin treatment on network synchronicity: firing rate (B), network burst rate (C), and percentage random spikes (D) in RHEB-p.P37L-expressing cultures. (E-F) Effect of vigabatrin treatment on network rhythmicity: NIBI (E) and coefficient of variance of NIBI (F) in RHEB-p.P37L-expressing cultures. (G-I) Effect of vigabatrin treatment on burst characteristics: network burst duration (G) network burst composition (H), and reverberation duration (I). Data is presented as mean ± SEM. Sample size (wells / replicates): CTR-VH: n = 24 / 2, CTR-VGB: n = 26 / 2, RHEB-p.P37L-VH: n = 26 / 2, RHEB-p.P37L-VGB: n = 24 / 2. Significant differences were determined using a Mixed-effect analysis with post hoc Bonferroni correction. * represent significant differences between vehicle (VH) and vigabatrin (VGB)-treated RHEB-p.P37L-expressing cultures. *p < 0.05, **p < 0.01, ***p < 0.001, ****p < 0.0001. All statistical comparisons are presented in Table 4A-B and S3.

Overall, vigabatrin initially reduced network activity to levels comparable to control cultures, but acted as a very strong suppressor of network activity in the following recording days. Interestingly, vigabatrin was able to introduce the organization of NBs into multiple reverberations, a pattern that is typically lost in mature neuronal cultures upon hyperactivity of the mTORC1 pathway.

## 3. DISCUSSION

In this study, we successfully identified reliable network activity parameters through the exploration of a variety of network activity parameters as outcome measures from the *in vitro* multiwell MEA system. We cultured mouse hippocampal neurons on the MEA and induced mTORC1 hyperactivity in two different ways: either through expression of the constitutively active RHEB-p.P37L variant, or knock-out of the *Tsc1* gene. The use of two different models to induce mTORC1-hyperactivity allowed us to identify shared phenotypes that depend on mTORC1-hyperactivity. We observed multiple changes in network dynamics, such as network synchronicity, network rhythmicity, and network burst characteristics (Table 3). More specifically, we identified increases in mean firing-and network burst rate and a faster decrease of the % random spikes. Additionally, we focused on identifying more robust outcome measures that were related to network activity. These included increased variability in the intervals between network bursts, and loss of the reverberating pattern in network bursts categorized as fewer reverberations per network burst and increased reverberation duration, which were consistently found to be affected throughout the experiments performed in this study.

Similar to previous studies in *Tsc1*-KO mouse (20) and TSC-patient iPSC-derived neuronal cultures (21–23), we observed increases in firing rate and network synchronization in mTORC1 hyperactive cultures. These parameters are consistently found to be increased throughout multiple studies, among which the phenotyping experiments in the current study. However, in our early rapamycin treatment experiments, we did not detect increases in firing rate and network synchronicity in the *Tsc1-*KO cultures. This supports the notion of a recent study using iPSC-derived neuron cultures, showing that these parameters are least robust and prone to technical and batch-to-batch variability (24). Therefore, we investigated the network behavior of our neuronal cultures in further detail. We first compared the rhythmicity of network activity and found that mTORC1 hyperactivity does consistently lead to more irregular activity patterns in mature neuronal cultures. Previously identified as robust outcome parameters (24), we subsequently investigated differences in network burst characteristics. Interestingly, we identified a drastic change in the network burst composition. Network bursts in mTORC1-hyperactive cultures generally consisted of single or very few reverberations of significantly prolonged duration compared to controls.

To investigate the mTOR dependency of the identified phenotypes, we treated cultures with the allosteric mTORC1 inhibitor rapamycin, starting either during early or late network development. We showed that early rapamycin treatment of mTORC1 hyperactive cultures could prevent the development of mTORC1-hyperactivity-induced network, and burst phenotypes. In contrast, late rapamycin treatment could not fully restore the identified phenotypes. Interestingly, while the mTORC1 hyperactive networks were unable to fire multiple reverberations in a single network burst upon late rapamycin treatment, it could reestablish a rhythmic busting pattern in the mTORC1 hyperactive cultures. We did not record the cultures after 3 weeks even though we did not yet see a rescue of all phenotypes when rapamycin treatment was started in the second week and could therefore not assess if prolonged exposure to rapamycin would eventually lead to a full rescue of the phenotypes. However, rapamycin acutely inhibits mTORC1 activity (31) and was shown to fully rescue epilepsy in *Tsc1-*KO and RHEB-p.P37L mouse models (26,32). Notably, network activity phenotypes were more profound in our RHEB-p.P37L cultures, compared to the *Tsc1*-KO model: phenotypes appeared earlier in development and differences were larger. This may indicate different pathological mechanisms but could more likely be a result of the method through which mTORC1 hyperactivity is induced. Cultures were manipulated using lentiviral transduction. While in our *Tsc1*^f/f^ cultures mTORC1 hyperactivity was induced through lentivirally expressed, Cre-mediated *Tsc1* knock out, RHEB-p.P37L was directly expressed from the lentiviral construct, resulting in a faster upregulation of the mTORC1 pathway. Furthermore, the direction in which parameters changed was the same in both mTORC1-hyperactivity models. On top of that, both models were sensitive to inhibition by the mTORC1-inhibitor rapamycin, suggesting that the network alterations primarily resulted from mTORC1-hyperactivity.

Patients with mTORopathy-related epilepsy often show poor responsiveness to AEDs. For example, a large cohort study showed that over 50% of TSC patients are prescribed three or more AEDs and another study showed that only 30% of patients attained seizure remission for longer than two years (10,33). This is a significant health burden for patients and their families and there is an urgent need to discover better AEDs. Therefore, we wanted to explore if the identified outcome parameters could be used as a screening assay for new AED treatments. To this extent, we assessed which of the identified phenotypes can be considered epileptic phenotypes, by treating the cultures with vigabatrin. Vigabatrin was previously shown to be the most effective in reducing seizure frequency in *Tsc1*-KO mice (32) and is the first line AED prescribed for infantile spasms in TSC patients (30). Vigabatrin enhances inhibitory gamma-aminobutyric acidergic (GABAergic) neurotransmission by inhibiting the breakdown of the neurotransmitter GABA by GABA transaminase (34). Within one hour, vigabatrin could dampen the heightened network activity and reduce the reverberation duration in RHEB-p.P37L cultures to levels similar to control cultures. We did not perform a dose titration study with vigabatrin and one day after the addition of vigabatrin, the drug exerted a strong suppressive effect on network activity of both control and RHEB-p.P37L cultures. Interestingly, even though activity levels were low in both vigabatrin-treated conditions on subsequent recording days, the drug was able to reintroduce the network burst composition in which multiple reverberations cluster together into a network burst in RHEB-p.P37L cultures, a phenotype that could not be restored by late rapamycin treatment.

The MEA system has previously been used to model epileptic phenotypes *in vitro* (15–17,35). During a specific period in brain development, highly synchronous, network-wide bursts are fired spontaneously. These spontaneous bursting events are actively involved in shaping neuronal networks (36–38). Although *in vitro* neuronal cultures have a simpler network organization, they also develop synchronous, network-wide bursting events which they maintain for the lifetime of the culture (39,40). *In vitro,* neuronal cultures may thus be used to study early network formation. The bursts that occur in neuronal cultures are shaped by an interplay between excitatory and inhibitory synapses (29,41,42). Previously, studies have shown that epileptic phenotypes could be introduced to neuronal networks on MEAs by pharmacological manipulations of the cultures affecting the balance between excitation and inhibition (15–17,35). The epileptic phenotypes that were identified in these studies included increased spike and network burst activity, as well as enhanced network synchronicity. Furthermore, these models were shown to be suitable to study the acute effects of epileptic drugs on network activity (35). In this study, we have shown that genetic manipulations that induce mTORC1-hyperactivity and cause epilepsy in patients, also alter network activity in neurons *in vitro* in a similar way as described by pharmacological manipulations as well as previous genetic mTORopathy models (20–23). Additionally, with our further assessment of network phenotypes in mTORC1 hyperactivity models, we expand the mTOR hyperactivity phenotypes in hippocampal neuronal cultures with alterations in network rhythmicity and burst characteristics. Interestingly, we observed a rescue of network burst reverberations when cultures were treated with vigabatrin, but this burst organization could not be restored when rapamycin was added to the cultures at similar time points. These data may suggest that mTORC1-hyperactivity early in development affects proper organization of the network into a balanced system that cannot be remodeled by inhibiting mTORC1-activity by rapamycin. However, targeting network excitability by enhancing GABAergic neurotransmission with vigabatrin could affect these parameters. While in *in* vivo animal models of mTOR-hyperactivity, rapamycin treatment can fully rescue epilepsy even after epilepsy onset (26,32), current clinical studies using rapalogs have not shown such strong benefits to patients. For example, a large phase 3 clinical study (EXIST-3) showed that the effectiveness of the mTORC1-inhibitor everolimus was considered clinically relevant in TSC patients with refractory epilepsy (i.e. ≥ 50% reduction in seizure frequency in 47% of patients) (43,44). However, patients did not reach seizure freedom. Currently, Vigabatrin is considered the first-line monotherapy for TSC-related epilepsy in young children (30). Retrospective and clinical trial studies with TSC patients have furthermore shown that prenatal diagnosis and preventive intervention with vigabatrin could reduce the incidence and severity of epilepsy and can improve TSC-associated neurological disorders (45–48). More recent findings however show contrasting results. The recent PREVeNT trial investigating the potential of preventative vigabatrin treatment, unfortunately, did not result in clinical improvement of epilepsy and behavioral outcomes (49). As of yet, it is still challenging to control seizures in mTORopahty patients, and a better understanding of disease mechanisms and development of new treatment strategies are needed.

Taken together, we have set up a valuable drug screening assay for mTORopathy-related epilepsies. We identified the multiwell MEA system as a useful model to study the development of neuronal activity during early network formation in a reduced system. Neuronal cultures can be manipulated easily and may thus also be used to further delineate which pathological processes in mTORopathies contribute to the alterations in network activity and to pathological mechanisms in the brains of mTORopathy patients. Another benefit of the multiwell MEA system is the reduced discomfort for experimental animals. While the effectiveness of new therapeutics should eventually always be tested *in vivo,* our *in vitro* system can provide an initial screening assay to select the most potent drugs for *in vivo* studies.

## 4. MATERIALS & METHODS

### 4.1 Mice

Mice were kept group-housed in IVC cages with bedding material on a 12/12-hour light/dark cycle at 21⁰C (±1⁰C), with a humidity level at 40-70%, and food pellets and water available ad libitum. When pregnant, mice were single housed. FvB/NHsD mice were used for RHEB-p.P37L experiments and *Tsc1*^f/f^ (*Tsc1*^tm1Djk^) mice were used for *Tsc1*-KO experiments (27). All animal experiments were conducted following the European Commission Council Directive 2010/63/EU (CCD project license AVD1010020172893 and AVD1010020172684), and protocols were subjected to ethical review (and approved) by an independent review board (IRB) of the Erasmus MC (studyplan numbers: SP2200020, SP2200045, and SP2200281).

### 4.2 Primary neuronal cultures

Primary hippocampal neuronal cultures were prepared according to the procedure described previously (50). For RHEBp.P37L-experiments, hippocampal neurons were isolated from FvB/NHsD embryos at embryonic day (E)16.5, and for *Tsc1*-KO experiments from *Tsc1*^f/f^ embryos at E17.5. Briefly, hippocampi were collected in Neurobasal medium (NBM, Gibco) on ice. Tissue was incubated for ∼20 minutes in pre-warmed trypsin/EDTA solution (Invitrogen) at 37⁰C. After washing with pre-warmed NB, cells were dissociated in NBM supplemented with 2% B27, 1% penicillin/streptomycin, and 1% glutamax (NBM^+++^), using a Pasteur pipette. For multi-electrode array experiments, 35,000 neurons were plated in a small drop in the center of each well in a 24-well multi-electrode array (MEA) plate, precoated with poly-d-lysine (25mg/ml) and airdried. Plates were then placed in an incubator at 37⁰C/5% CO_2_ for 45 minutes to ensure attachments of neurons on top of the electrodes. Finally, 500 uL pre-warmed, NBM^+++^ was added to the wells. For western blot (WB) experiments, 300,000 neurons/well were plated in NBM^+++^, in a 12-well plate precoated with poly-d-lysine.

### 4.3 RHEB construct

cDNA encoding the *RHEB* (NM_005614.3) c.110C*>*T (p.P37L) mutation was synthesized by GeneCust as described previously (25,26). This was cloned as an AscI/PacI fragment into the multiple cloning site downstream of the EF1a promoter in the pLVX-EF1α-IRES-ZsGreen1 Vector (Takara). The WT allele was generated by site-directed mutagenesis using the following 2 primers (Forward 5’-atttgtggactcctacgatcCaaccatagaaaacactttta-3’ and Reverse 5’-taaaagtgttttctatggttGgatcgtaggagtccacaaat-3’.

### 4.4 Lentiviral production

HEK293T cells were cultured in DMEM supplemented with 10% Fetal Bovine Serum and 1% penicillin/streptomycin and incubated for 5 hours with Chloroquine diphosphate (25 μM, C6628-25G, Sigma-Aldrich) added to the medium. A DNA mix containing the required plasmids (pMD2.G (#12259 Addgene), psPAX2 (#12260 Addgene), and the empty (EV), RHEB-WT or RHEB-p.P37L expressing lentiviral plasmid) and polyethylenimine (DNA:PEI ratio 1:3) was incubated for 15-20 minutes at room temperature, subsequently added dropwise to the HEK293T cells and incubated overnight. The medium was refreshed the next day and virus-containing medium was collected and concentrated 72 hours post-refreshing using Amicon Ultra 15 filters (C7715, Merck Millipore).

### 4.5 Lentiviral transduction

Primary hippocampal neuronal cultures were transduced with lentivirus (LV) at 1 day in vitro (DIV1). For the induction of *Tsc1*-gene deletion, 1μL or 2 μL (for MEA and WB experiments respectively) of 1*10^7^ CFU/mL of a commercial PGK-cre-GFP virus (Cellomics Technology, PLV-10079) was used. As a control, wells were transduced with 1 μL (MEA) or 2 μL (WB) LV-EV. For RHEB experiments, wells were transduced with 0.5 μL of either LV-EV, LV-RHEB-WT virus or LV-RHEB-p.P37L virus for MEA experiments and with 1 uL of each lentivirus for WB experiments.

### 4.6 Western blot analysis

At DIV14, cells were lysed with a sucrose buffer consisting of 250 mM sucrose, 20 mM Hepes pH 7.2, 1 mM MgCl_2_, and 10 U/mL Benzonase, in the presence of phosphatase and proteinase inhibitors. Protein concentration was determined using a BCA protein assay kit (Thermo Fisher Scientific, 23225). Approximately 15 μg protein was separated by SDS-PAGE on a precast 4-12% Criterion XT Bis-Tris gel (Bio-Rad) and transferred to a nitrocellulose membrane using TurboBlot (Bio-Rad). Membranes were blocked for 1 hour at room temperature in 5% powdered milk dissolved in TBS-T: Tris-buffered saline (10mM Tris-HCl, pH 8.0, 150 mM NaCl) with 0.1% Tween-20 (Sigma P1379). Blots were incubated with the primary antibodies diluted in TBS-T overnight at 4°C. Blots were washed with TBS-T 3 times for 10 minutes and incubated with secondary antibodies diluted in TBS-T for 1 hour at room temperature. Membranes were then washed 2 times for 10 minutes in TBS-T and a final wash for 10 minutes in TBS. The following primary antibodies were used: rabbit anti-Phospho-S6 Ribosomal Protein (Ser240/244) (1:2000, #5364 Cell Signaling), rabbit anti-S6 Ribosomal Protein (1:2000 #2217 Cell Signaling), rabbit anti-TSC1 (1:1000, #6935 Cell Signaling) and mouse anti-Glyceraldehyde-3-Phosphate Dehydrogenase (GAPDH) (1:10,000, MAB374, Sigma-Aldrich). Secondary antibodies used were: goat anti-rabbit (680 nm) and goat anti-mouse (800 nm) conjugates (1:15,000, Li-Cor Biosciences, Lincoln, USA). Fluorescence was detected on an Odyssey near-infrared scanner and protein levels were quantified using the Image Studio Lite version 5.2 software.

### 4.7 MEA recordings

24-well MEA plates with an epoxy base (Multichannel Systems, MCS GmbH, Reutlingen, Germany) were used to record spontaneous network activity. Each well has embedded a 4×4 grid of PEDOT-coated gold electrodes with a diameter of 100 μm, each spaced 700 μm apart. Plates were recorded using the multiwell-MEA headstage in a recording chamber at 37⁰C / 5% CO_2_. After placement of the plate into the recording headstage, cells were left to acclimatize for 10 minutes. Subsequently, activity was recorded for 10 minutes with a sampling rate of 10 kHz. The signal was filtered with a 4^th^-order low-pass filter at 3.5 kHz and 2^nd^-order high-pass filter at 100 Hz. For RHEB experiments, network activity was recorded at DIVs 5, 7, 10, 14, 16, and 19. For *Tsc1*-KO experiments, network activity was recorded at DIVs 4, 7, 11, 14, 18, and 21. Recordings for vigabatrin experiments were performed at DIVs 5, 7, 9, 11 (1x pre-and 3x post-treatment), 12, 14, 16 and 19. After every recording (except DIV11 1x pre-and 3x post-treatment recordings in vigabatrin experiments), one-third of the medium was replaced with fresh NBM^+++^.

### 4.8 Drug treatment

Rapamycin was dissolved in dimethyl sulfoxide (DMSO) and added to the cultures after recording at DIV3 or DIV10 (RHEB experiments), or DIV4 or DIV14 (*Tsc1*-KO experiments) in a concentration of 50 nM rapamycin with DMSO diluted 1:4000. As a control, cultures were treated with 1:4000 DMSO only. Drug concentration was maintained with every refreshing step. Treatment of western blot cultures was done 4 days before harvesting the cells. Vigabatrin was dissolved in sterile H_2_O and added to the cultures at DIV11 with a final concentration of 25 µM, by a 1:500 dilution. As a control, cultures were treated with 1:500 dilution of sterile H_2_O only. After the recording at DIV12, vigabatrin was gradually washed out by using drug-free NBM^+++^ for refreshing.

### 4.9 MEA data analysis and statistics

Data analysis was performed offline using the Multiwell Analyzer software (Multichannel Systems, MCS GmbH, Reutlingen, Germany). Analysis was performed on the last 5 minutes of the recording period. Baseline noise was calculated as the average of 2x 200ms segments without activity. Spikes were detected using a threshold of +/-5 standard deviations from baseline. Reverberations were detected in the Multiwell Analyzer software using the parameters in Table 5.

**Table 5.**
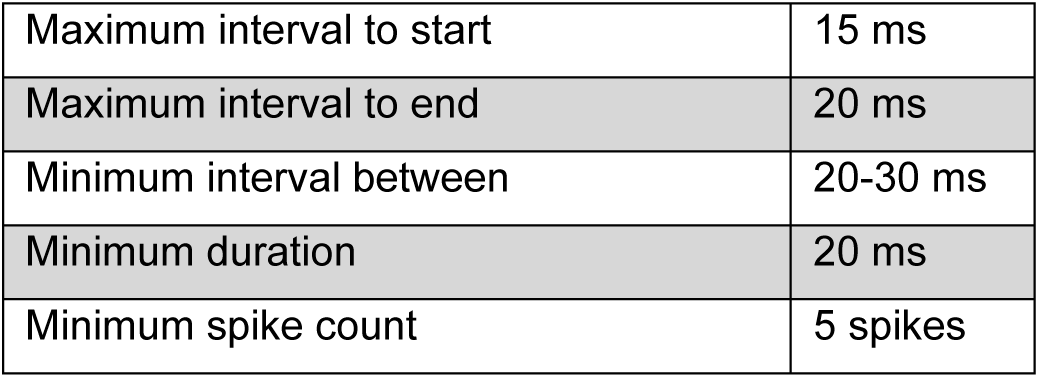
Reverberation definition parameters.

Network reverberations were defined as reverberations in which at least two-thirds of the channels in one well were participating, of which half of them were active simultaneously. After spike-and reverberation detection were performed in the MSC software, data was further processed using MATLAB R2021a (The MathWorks, Inc., Natick, Massachusetts, United States). Reverberations were combined into network bursts when the interval to the next reverberation was < 300 ms. Occasionally, in cultures of at least 2 weeks old, wells showed very little, to no activity at all at a single DIV, while activity levels returned to normal on the next recording day (for an example see Fig S2). Since activity is required for the characterization of network activity changes, silent recording days, defined as recordings (DIV14 and older) with a network burst rate ≤ 0.5 NB/min, were excluded from the dataset. In the vigabatrin experiments, wells with a network burst rate ≤ 0.5 NB/min were included as this was likely due to the strong suppressive action of the drug on network activity. In this case, outcome parameters that could not be calculated in the absence of bursts (network inter-burst interval (NIBI), coefficient of variance of NIBI (CoV^NIBI^), network burst duration (NBD), network burst composition (NBC) and reverberation duration) were reported as NaN. Statistical analysis was performed using a Student *t-*test, One-way ANOVA, or Mixed model analysis, corrected for multiple comparisons using Bonferroni correction. For all statistical analyses, α was set at 0.05. The specific tests used for each experiment are specified in the figure legends or the results section. Values are represented as averages ± SEM. Sample sizes for each experiment are indicated in the figure legends. Statistical analyses were performed using GraphPad Prism 5 (GraphPad Software, Inc., CA, USA).

## ACKNOWLEDGEMENT

The authors would like to thank Charlotte de Konink for her technical contributions and management of the mouse colony.

## Funding

This work was supported by the Dutch TSC Foundation (STSN) (G.M.v.W) and the Dutch Epilepsy Foundation (Epilepsie fonds) (Y.E.). GMvW was supported by the Dutch Research Council (NWO) Vidi Grant (016.Vidi.188014). The funders had no role in study design, data collection and analysis, decision to publish, or preparation of the manuscript.

## AUTHOR CONTRIBUTIONS

AMH, MPO, YE, and GvW: Study design and interpretation; AMH, MPO, JDdH, and EN: Data collection and analysis; MF: methodological support; AMH: writing manuscript; all authors: manuscript review.

## CONFLICT OF INTEREST

MPO is currently working as a senior scientist at Ionis.

