## Supplementary material for "Modeling mTORopathy-related epilepsy in cultured murine hippocampal neurons using the multi-electrode array"

### SUPPLEMENTARY FIGURE 1

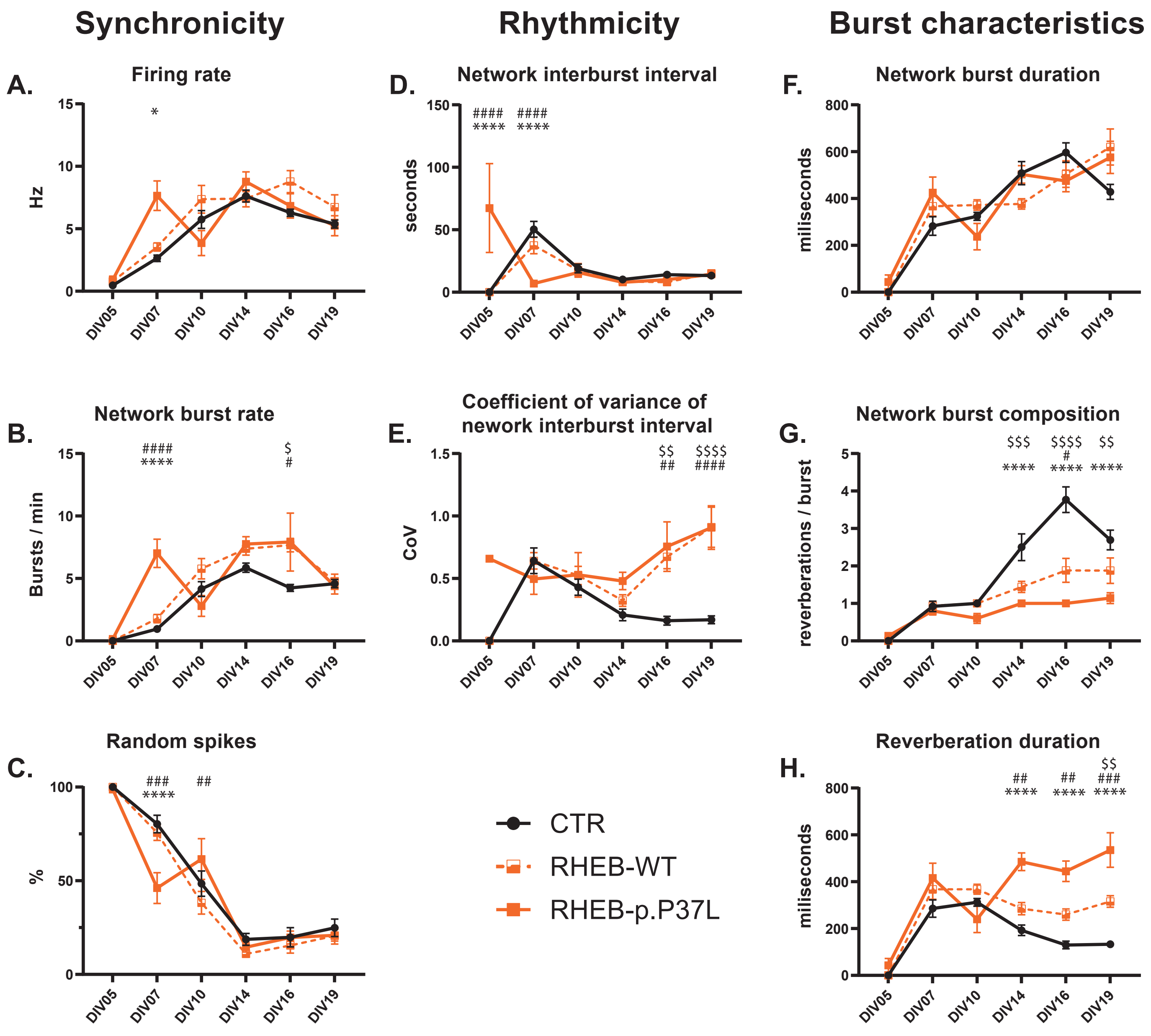

### SUPPLEMENTARY FIGURE 2

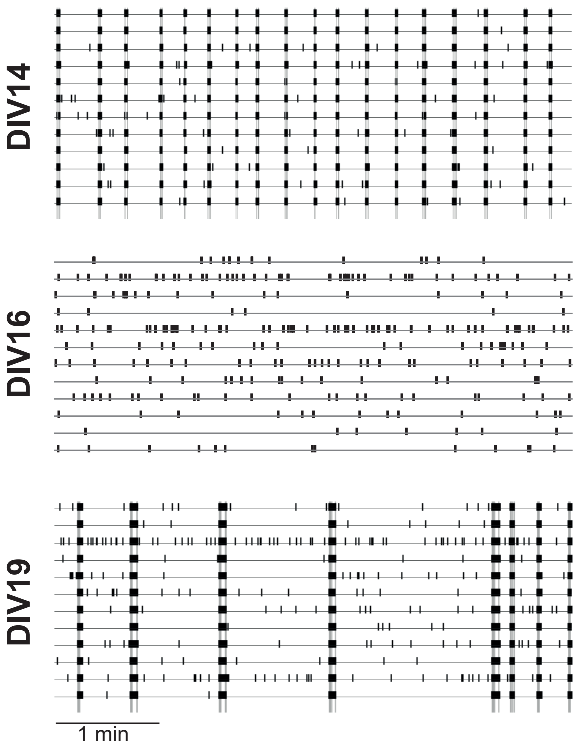

#### SUPPORTING INFORMATION

<u>S1 fig. Overexpression of RHEB-WT causes intermediate phenotypes on the MEA.</u> Cultures transduced with RHEB-WT show less severe phenotypes than cultures transduced with RHEB-p.P37L. (A) firing rate, (B) network burst rate, (C) percentage random spikes, (D) NIBI, (E) coefficient of variance of NIBI, (F) network burst duration, (G) network burst composition, and (H) reverberation duration. Data is presented as mean ± SEM. Sample size (wells/replicates): CTR: n = 13 / 3, RHEB-p.P37L: n = 17 / 3, RHEB-WT: n = 15 / 3. Significant differences were determined using a Mixed-effect analysis with post hoc Bonferroni correction. * represent significant differences between RHEB-p.P37L and CTR, # represent significant differences between RHEB-p.P37L and RHEB-WT, $ represent significant differences between RHEB-WT and CTR. *p < 0.05, **p < 0.01, ***p < 0.001, ****p < 0.0001. All statistical comparisons are presented in Supplementary table 1A-B.

<u>S2 fig. Example raster plot of silent wells at DIV16/18 that were excluded from analysis.</u> Example of a 12-electrode raster plot on DIV14, 16 and 19. Each horizontal line represents a single electrode, with vertical lines representing spikes. Gray bars spanning the entire raster represent network bursts. Well shows expected network activity on DIV14 and DIV19 but no network activity, only random spikes, on DIV16.

**S1A table. Statistical analysis overview of mixed effects analysis for establishment of RHEB-WT and RHEB-p.P37L induced phenotypes on the MEA.**

| Outcome parameter | Between effect | | Interaction effect | |
|---|---|---|---|---|
| | **p-value** | **F** | **p-value** | **F** |
| **MFR** | 0.2023 | (2, 42) = 1.660 | ****<0.0001 | (10, 206) = 5.340 |
| **%RS** | 0.2322 | (2, 42) = 1.512 | ****<0.0001 | (10, 206) = 4.186 |
| **NBR** | **0.0049 | (2, 42) = 6.057 | ****<0.0001 | (10, 206) = 5.102 |
| **NIBI** | *0.0290 | (2, 42) = 3.855 | ****<0.0001 | (10, 177) = 14.96 |
| **$CoV^{NIBI}$** | ****<0.0001 | (1, 219) = 14.55 | ***0.0003 | (10, 219) = 3.516 |
| **NBD** | 0.7583 | (2, 42) = 0.2785 | **0.0021 | (10, 205) = 2.900 |
| **NBC** | ****<0.0001 | (2, 42) = 29.12 | ****<0.0001 | (10, 206) = 9.542 |
| **Reverberation duration** | ****<0.0001 | (2, 248) = 36.54 | ****<0.0001 | (10, 248) = 6.703 |

**MFR = Mean firing rate, %RS = percentage random spikes, NIBI = network inter-burst interval, $CoV^{NIBI}$ = Coefficient of variance of NIBI, NBR = Network burst rate, NBD = Network burst duration, NBC = Network burst composition.**

#### S1B table. Post-hoc Bonferroni corrected multiple comparisons for the establishment of RHEB-induced phenotypes on the MEA.

| Parameter | Comparison | p-value | | | | | |
|---|---|---|---|---|---|---|---|
| | | **DIV05** | **DIV07** | **DIV10** | **DIV14** | **DIV18** | **DIV19/21** |
| **MFR** | CTR – RHEB-WT | 0.9976 | 0.7035 | >0.9999 | >0.9999 | 0.2569 | >0.9999 |
| | CTR – RHEB-p.P37L | >0.9999 | **0.0151** | >0.9999 | >0.9999 | >0.9999 | >0.9999 |
| | RHEB-WT – RHEB-p.P37L | >0.9999 | 0.0725 | 0.4579 | >0.9999 | >0.9999 | >0.9999 |
| **%RS** | CTR – RHEB-WT | >0.9999 | >0.9999 | >0.9999 | >0.9999 | >0.9999 | >0.9999 |
| | CTR – RHEB-p.P37L | >0.9999 | **<0.0001** | >0.9999 | >0.9999 | >0.9999 | >0.9999 |
| | RHEB-WT – RHEB-p.P37L | >0.9999 | **0.0003** | **0.0102** | >0.9999 | >0.9999 | >0.9999 |
| **NBR** | CTR – RHEB-WT | >0.9999 | >0.9999 | >0.9999 | >0.9999 | **0.0264** | >0.9999 |
| | CTR – RHEB-p.P37L | >0.9999 | **<0.0001** | >0.9999 | >0.9999 | **0.0200** | >0.9999 |
| | RHEB-WT – RHEB-p.P37L | >0.9999 | **<0.0001** | 0.0713 | >0.9999 | >0.9999 | >0.9999 |
| **NIBI** | CTR – RHEB-WT | >0.9999 | 0.1542 | >0.9999 | >0.9999 | >0.9999 | >0.9999 |
| | CTR – RHEB-p.P37L | **<0.0001** | **<0.0001** | >0.9999 | >0.9999 | >0.9999 | >0.9999 |
| | RHEB-WT – RHEB-p.P37L | **<0.0001** | **<0.0001** | >0.9999 | >0.9999 | >0.9999 | >0.9999 |
| **$CoV^{NIBI}$** | CTR – RHEB-WT | >0.9999 | >0.9999 | >0.9999 | >0.9999 | **0.0054** | **<0.0001** |
| | CTR – RHEB-p.P37L | 0.4146 | >0.9999 | >0.9999 | >0.9999 | **0.0012** | **<0.0001** |
| | RHEB-WT – RHEB-p.P37L | 0.3770 | >0.9999 | >0.9999 | >0.9999 | >0.9999 | >0.9999 |
| **NBD** | CTR – RHEB-WT | >0.9999 | >0.9999 | >0.9999 | 0.7221 | >0.9999 | 0.0614 |
| | CTR – RHEB-p.P37L | >0.9999 | 0.6185 | >0.9999 | >0.9999 | >0.9999 | 0.511 |
| | RHEB-WT – RHEB-p.P37L | >0.9999 | >0.9999 | 0.5066 | 0.7851 | >0.9999 | >0.9999 |
| **NBC** | CTR – RHEB-WT | >0.9999 | >0.9999 | >0.9999 | **0.0015** | **<0.0001** | **0.0405** |
| | CTR – RHEB-p.P37L | >0.9999 | >0.9999 | >0.9999 | **<0.0001** | **<0.0001** | **<0.0001** |
| | RHEB-WT – RHEB-p.P37L | >0.9999 | >0.9999 | >0.9999 | >0.9999 | **0.0142** | 0.1179 |
| **Reverberation duration** | CTR – RHEB-WT | >0.9999 | >0.9999 | >0.9999 | >0.9999 | 0.2156 | **0.0093** |
| | CTR – RHEB-p.P37L | >0.9999 | 0.2576 | >0.9999 | **<0.0001** | **<0.0001** | **<0.0001** |
| | RHEB-WT – RHEB-p.P37L | >0.9999 | >0.9999 | 0.1771 | **0.0015** | **0.0049** | **0.0004** |

**MFR = Mean firing rate, %RS = percentage random spikes, NIBI = network inter-burst interval, $CoV^{NIBI}$ = Coefficient of variance of NIBI, NBR = Network burst rate, NBD = Network burst duration, NBC = Network burst composition.**

#### S2A table. Post-hoc Bonferroni corrected multiple comparisons for early and late rescue of RHEB-p.P37L induced mTORC1 hyperactivity phenotypes on the MEA.

| Parameter | Comparison | p-value | | | | | |
|---|---|---|---|---|---|---|---|
| | | **DIV04/05** | **DIV07** | **DIV10/11** | **DIV14** | **DIV16/18** | **DIV19/21** |
| **Early treatment – RHEB-p.P37L** | | | | | | | |
| **MFR** | CTR-DMSO – RHEB-p.P37L-DMSO | **0.0203** | **0.0005** | **0.0014** | >0.9999 | 0.4525 | >0.9999 |
| | CTR-RAPA – RHEB-p.P37L-DMSO | **0.0005** | **0.0002** | **0.0083** | >0.9999 | 0.6786 | >0.9999 |
| | CTR-RAPA – RHEB-p.P37L-RAPA | 0.2898 | >0.9999 | 0.3800 | >0.9999 | 0.9727 | >0.9999 |
| **%RS** | CTR-DMSO – RHEB-p.P37L-DMSO | >0.9999 | **0.0098** | **<0.0001** | >0.9999 | >0.9999 | >0.9999 |
| | CTR-RAPA – RHEB-p.P37L-DMSO | >0.9999 | **0.0004** | **0.0002** | >0.9999 | 0.2114 | 0.8689 |
| | CTR-RAPA – RHEB-p.P37L-RAPA | | 0.7342 | >0.9999 | >0.9999 | **0.0322** | >0.9999 |
| **NBR** | CTR-DMSO – RHEB-p.P37L-DMSO | >0.9999 | **0.0018** | **0.0059** | **0.0023** | **0.0409** | 0.2269 |
| | CTR-RAPA – RHEB-p.P37L-DMSO | >0.9999 | **0.0004** | **0.0156** | 0.0657 | **0.0190** | >0.9999 |
| | CTR-RAPA – RHEB-p.P37L-RAPA | | >0.9999 | >0.9999 | >0.9999 | 0.3447 | >0.9999 |
| **NIBI** | CTR-DMSO – RHEB-p.P37L-DMSO | | 0.3030 | **0.0001** | **0.0026** | >0.9999 | **0.0326** |
| | CTR-RAPA – RHEB-p.P37L-DMSO | | | **0.0038** | **0.0082** | >0.9999 | 0.7669 |

| | | | | | | | |
|---|---|---|---|---|---|---|---|
| | CTR-RAPA – RHEB-p.P37L-RAPA | | | >0.9999 | >0.9999 | 0.7068 | >0.9999 |
| **$CoV^{NIBI}$** | CTR-DMSO – RHEB-p.P37L-DMSO | | 0.4539 | **0.0004** | **0.0079** | **0.0002** | **0.0002** |
| | CTR-RAPA – RHEB-p.P37L-DMSO | | | **0.0012** | **0.0236** | >0.9999 | 0.2299 |
| | CTR-RAPA – RHEB-p.P37L-RAPA | | | 0.8944 | >0.9999 | **0.0248** | 0.0502 |
| **NBD** | CTR-DMSO – RHEB-p.P37L-DMSO | >0.9999 | 0.0713 | 0.1362 | 0.1728 | >0.9999 | >0.9999 |
| | CTR-RAPA – RHEB-p.P37L-DMSO | >0.9999 | **<0.0001** | 0.0527 | 0.7779 | >0.9999 | >0.9999 |
| | CTR-RAPA – RHEB-p.P37L-RAPA | | >0.9999 | >0.9999 | >0.9999 | 0.9050 | **0.0315** |
| **NBC** | CTR-DMSO – RHEB-p.P37L-DMSO | >0.9999 | >0.9999 | >0.9999 | **<0.0001** | **<0.0001** | **<0.0001** |
| | CTR-RAPA – RHEB-p.P37L-DMSO | >0.9999 | **0.0096** | >0.9999 | **0.0013** | **0.0014** | 0.0988 |
| | CTR-RAPA – RHEB-p.P37L-RAPA | | >0.9999 | >0.9999 | >0.9999 | 0.7812 | 0.0823 |
| **Reverberation duration** | CTR-DMSO – RHEB-p.P37L-DMSO | >0.9999 | 0.0595 | 0.0955 | **<0.0001** | **0.0001** | **0.0001** |
| | CTR-RAPA – RHEB-p.P37L-DMSO | >0.9999 | **<0.0001** | 0.0572 | **0.0002** | **0.0002** | **0.0002** |
| | CTR-RAPA – RHEB-p.P37L-RAPA | | >0.9999 | >0.9999 | >0.9999 | >0.9999 | >0.9999 |
| | | **DIV04/05** | **DIV07** | **DIV10/11** | **DIV14** | **DIV16/18** | **DIV19/21** |
| **Late treatment – RHEB-p.P37L** | | | | | | | |
| **MFR** | CTR-DMSO – RHEB-p.P37L-DMSO | **0.0015** | **<0.0001** | **0.0041** | >0.9999 | 0.8610 | >0.9999 |
| | CTR-RAPA – RHEB-p.P37L-DMSO | **0.0025** | **<0.0001** | **0.0487** | >0.9999 | 0.2730 | 0.6137 |
| | CTR-RAPA – RHEB-p.P37L-RAPA | **0.0099** | **0.0007** | >0.9999 | **0.0024** | >0.9999 | >0.9999 |
| **%RS** | CTR-DMSO – RHEB-p.P37L-DMSO | 0.1376 | **<0.0001** | **0.0005** | 0.3048 | >0.9999 | >0.9999 |
| | CTR-RAPA – RHEB-p.P37L-DMSO | 0.1376 | **<0.0001** | **0.0179** | >0.9999 | 0.0501 | **0.0354** |
| | CTR-RAPA – RHEB-p.P37L-RAPA | 0.1733 | **0.0023** | **0.0448** | **0.0163** | >0.9999 | >0.9999 |
| **NBR** | CTR-DMSO – RHEB-p.P37L-DMSO | 0.1857 | **<0.0001** | **<0.0001** | 0.0989 | 0.0564 | >0.9999 |
| | CTR-RAPA – RHEB-p.P37L-DMSO | 0.1857 | **<0.0001** | **0.0005** | 0.4532 | **0.0011** | 0.5777 |
| | CTR-RAPA – RHEB-p.P37L-RAPA | 0.2491 | **0.0003** | **0.0230** | 0.1176 | **0.0134** | >0.9999 |
| **NIBI** | CTR-DMSO – RHEB-p.P37L-DMSO | **0.0367** | **<0.0001** | **<0.0001** | **0.0389** | >0.9999 | >0.9999 |
| | CTR-RAPA – RHEB-p.P37L-DMSO | **0.0367** | **<0.0001** | **0.0004** | **0.0303** | 0.2569 | **0.0400** |
| | CTR-RAPA – RHEB-p.P37L-RAPA | 0.0780 | 0.1878 | **0.0379** | **0.0718** | **0.0228** | >0.9999 |
| **$CoV^{NIBI}$** | CTR-DMSO – RHEB-p.P37L-DMSO | **0.0015** | **0.0364** | **<0.0001** | 0.1064 | **0.0101** | **0.0047** |
| | CTR-RAPA – RHEB-p.P37L-DMSO | **0.0015** | 0.0503 | **0.0020** | **0.0166** | 0.2910 | 0.3494 |
| | CTR-RAPA – RHEB-p.P37L-RAPA | **0.0031** | 0.0728 | **0.0004** | 0.0877 | >0.9999 | 0.1593 |
| **NBD** | CTR-DMSO – RHEB-p.P37L-DMSO | 0.0682 | **0.0004** | >0.9999 | 0.0833 | 0.175 | 0.8574 |
| | CTR-RAPA – RHEB-p.P37L-DMSO | 0.0682 | **<0.0001** | >0.9999 | **0.0223** | 0.0588 | >0.9999 |
| | CTR-RAPA – RHEB-p.P37L-RAPA | **0.0406** | **0.0188** | >0.9999 | **0.0083** | **<0.0001** | **0.0009** |
| **NBC** | CTR-DMSO – RHEB-p.P37L-DMSO | 0.0595 | >0.9999 | 0.9770 | **<0.0001** | **<0.0001** | 0.3769 |
| | CTR-RAPA – RHEB-p.P37L-DMSO | 0.0595 | 0.1274 | 0.9770 | **<0.0001** | **<0.0001** | 0.3019 |
| | CTR-RAPA – RHEB-p.P37L-RAPA | **0.0406** | 0.1274 | >0.9999 | **0.0037** | **<0.0001** | **0.0265** |
| **Reverberation duration** | CTR-DMSO – RHEB-p.P37L-DMSO | 0.0682 | **0.0005** | >0.9999 | **0.0086** | **<0.0001** | **<0.0001** |
| | CTR-RAPA – RHEB-p.P37L-DMSO | 0.0682 | **<0.0001** | >0.9999 | **0.0066** | **<0.0001** | **0.0001** |
| | CTR-RAPA – RHEB-p.P37L-RAPA | **0.0423** | **0.0156** | 0.1871 | >0.9999 | **0.0001** | **0.0108** |

**MFR = Mean firing rate, %RS = percentage random spikes, NIBI = network inter-burst interval, $CoV^{NIBI}$ = Coefficient of variance of NIBI, NBR = Network burst rate, NBD = Network burst duration, NBC = Network burst composition.**

#### S2B table. Post-hoc Bonferroni corrected multiple comparisons for early and late rescue of *Tsc1*-KO induced mTORC1 hyperactivity phenotypes on the MEA.

| Parameter | Comparison | p-value | | | | | |
|---|---|---|---|---|---|---|---|
| | | **DIV04/05** | **DIV07** | **DIV10/11** | **DIV14** | **DIV16/18** | **DIV19/21** |
| **Early treatment – *Tsc1*-KO** | | | | | | | |
| **MFR** | CTR-DMSO – CRE-DMSO | >0.9999 | >0.9999 | >0.9999 | 0.2691 | 0.1001 | >0.9999 |
| | CTR-RAPA – CRE-DMSO | >0.9999 | >0.9999 | **0.0069** | 0.2171 | 0.3921 | 0.2490 |
| | CTR-RAPA – CRE-RAPA | 0.2105 | >0.9999 | >0.9999 | 0.1308 | **0.0459** | 0.1602 |
| **%RS** | CTR-DMSO – CRE-DMSO | | **0.0136** | >0.9999 | 0.2184 | 0.4341 | >0.9999 |
| | CTR-RAPA – CRE-DMSO | | >0.9999 | **0.0064** | **0.0178** | 0.3464 | >0.9999 |
| | CTR-RAPA – CRE-RAPA | | >0.9999 | >0.9999 | >0.9999 | **0.0046** | >0.9999 |
| **NBR** | CTR-DMSO – CRE-DMSO | | 0.4957 | >0.9999 | 0.4086 | 0.1177 | >0.9999 |
| | CTR-RAPA – CRE-DMSO | | >0.9999 | **0.0274** | 0.0778 | 0.0571 | 0.2010 |
| | CTR-RAPA – CRE-RAPA | | >0.9999 | >0.9999 | >0.9999 | 0.0661 | >0.9999 |

| | | | | | | | |
|---|---|---|---|---|---|---|---|
| **NIBI** | CTR-DMSO – CRE-DMSO | | >0.9999 | >0.9999 | >0.9999 | 0.1682 | >0.9999 |
| | CTR-RAPA – CRE-DMSO | | >0.9999 | **0.0057** | 0.0846 | 0.1570 | >0.9999 |
| | CTR-RAPA – CRE-RAPA | | 0.9345 | >0.9999 | 0.7081 | 0.0742 | >0.9999 |
| **CoV$^{NIBI}$** | CTR-DMSO – CRE-DMSO | | >0.9999 | >0.9999 | >0.9999 | >0.9999 | 0.2077 |
| | CTR-RAPA – CRE-DMSO | | >0.9999 | >0.9999 | 0.1311 | 0.1719 | >0.9999 |
| | CTR-RAPA – CRE-RAPA | | >0.9999 | >0.9999 | >0.9999 | >0.9999 | 0.2975 |
| **NBD** | CTR-DMSO – CRE-DMSO | | 0.2540 | >0.9999 | 0.4877 | >0.9999 | >0.9999 |
| | CTR-RAPA – CRE-DMSO | | >0.9999 | **0.0026** | >0.9999 | >0.9999 | >0.9999 |
| | CTR-RAPA – CRE-RAPA | | >0.9999 | 0.0547 | >0.9999 | 0.1763 | >0.9999 |
| **NBC** | CTR-DMSO – CRE-DMSO | | 0.1025 | 0.2602 | **0.0050** | 0.3326 | 0.4549 |
| | CTR-RAPA – CRE-DMSO | | >0.9999 | 0.8070 | 0.1194 | **<0.0001** | **0.0373** |
| | CTR-RAPA – CRE-RAPA | | >0.9999 | >0.9999 | >0.9999 | **0.0025** | 0.1588 |
| **Reverberation duration** | CTR-DMSO – CRE-DMSO | | 0.2235 | **0.0007** | **<0.0001** | **0.0096** | **0.0448** |
| | CTR-RAPA – CRE-DMSO | | >0.9999 | **<0.0001** | **<0.0001** | **0.0005** | 0.0771 |
| | CTR-RAPA – CRE-RAPA | | >0.9999 | >0.9999 | >0.9999 | 0.2209 | 0.9536 |
| | | **DIV04/05** | **DIV07** | **DIV10/11** | **DIV14** | **DIV16/18** | **DIV19/21** |
| **Late treatment – *Tsc1*-KO** | | | | | | | |
| **MFR** | CTR-DMSO – CRE-DMSO | 0.4572 | **0.0331** | >0.9999 | >0.9999 | >0.9999 | >0.9999 |
| | CTR-RAPA – CRE-DMSO | 0.3061 | **0.0020** | >0.9999 | >0.9999 | 0.5343 | >0.9999 |
| | CTR-RAPA – CRE-RAPA | >0.9999 | **0.0071** | 0.1177 | >0.9999 | >0.9999 | 0.0532 |
| **%RS** | CTR-DMSO – CRE-DMSO | | 0.0891 | >0.9999 | >0.9999 | >0.9999 | >0.9999 |
| | CTR-RAPA – CRE-DMSO | | 0.2253 | >0.9999 | >0.9999 | >0.9999 | 0.4934 |
| | CTR-RAPA – CRE-RAPA | | >0.9999 | 0.2066 | 0.6206 | 0.8528 | **0.0407** |
| **NBR** | CTR-DMSO – CRE-DMSO | | **0.0156** | >0.9999 | 0.0717 | >0.9999 | >0.9999 |
| | CTR-RAPA – CRE-DMSO | | **0.0154** | 0.3348 | **0.032** | >0.9999 | >0.9999 |
| | CTR-RAPA – CRE-RAPA | | 0.1371 | **0.0099** | **<0.0001** | >0.9999 | 0.6225 |
| **NIBI** | CTR-DMSO – CRE-DMSO | | 0.7793 | >0.9999 | >0.9999 | >0.9999 | >0.9999 |
| | CTR-RAPA – CRE-DMSO | | >0.9999 | 0.1004 | 0.9997 | >0.9999 | >0.9999 |
| | CTR-RAPA – CRE-RAPA | | >0.9999 | **0.0129** | **0.0005** | >0.9999 | >0.9999 |
| **CoV$^{NIBI}$** | CTR-DMSO – CRE-DMSO | | >0.9999 | >0.9999 | 0.9026 | >0.9999 | 0.8381 |
| | CTR-RAPA – CRE-DMSO | | **0.0156** | >0.9999 | >0.9999 | >0.9999 | >0.9999 |
| | CTR-RAPA – CRE-RAPA | | **0.0269** | 0.1641 | 0.5670 | 0.2878 | 0.1300 |
| **NBD** | CTR-DMSO – CRE-DMSO | | 0.1870 | 0.2037 | 0.2372 | >0.9999 | 0.7377 |
| | CTR-RAPA – CRE-DMSO | | 0.1568 | >0.9999 | **0.0168** | 0.1561 | 0.8809 |
| | CTR-RAPA – CRE-RAPA | | **0.0005** | >0.9999 | **0.002** | **0.0328** | **0.0024** |
| **NBC** | CTR-DMSO – CRE-DMSO | | **0.0474** | 0.4937 | **0.0487** | >0.9999 | **0.0052** |
| | CTR-RAPA – CRE-DMSO | | **0.0131** | 0.9227 | **0.0277** | **0.0008** | **0.0003** |
| | CTR-RAPA – CRE-RAPA | | **0.0002** | 0.9227 | **0.0239** | **0.0011** | **0.0174** |
| **Reverberation duration** | CTR-DMSO – CRE-DMSO | | 0.1454 | 0.1409 | **0.0005** | >0.9999 | **0.0009** |
| | CTR-RAPA – CRE-DMSO | | 0.1405 | **0.0055** | **<0.0001** | **0.0443** | **0.0006** |
| | CTR-RAPA – CRE-RAPA | | **0.0006** | 0.2897 | **0.0003** | **<0.0001** | **0.0183** |

**MFR = Mean firing rate, %RS = percentage random spikes, NIBI = network inter-burst interval, CoV$^{NIBI}$ = Coefficient of variance of NIBI, NBR = Network burst rate, NBD = Network burst duration, NBC = Network burst composition.**

**S3 table. Post-hoc Bonferroni corrected multiple comparisons for vigabatrin treatment in RHEB-p.P37L-induced mTORC1 hyperactivity phenotypes on the MEA.**

| Parameter | Comparison | p-value | | | | | | | | | | |
|---|---|---|---|---|---|---|---|---|---|---|---|---|
| | | **DIV05** | **DIV07** | **DIV09** | **DIV11** | **DIV11 + 1hr** | **DIV11 + 3hr** | **DIV11 + 5hr** | **DIV12** | **DIV14** | **DIV16** | **DIV19** |
| **MFR** | CTR-VH – RHEB-p.P37L-VH | **0.0477** | **<0.0001** | **<0.0001** | **<0.0001** | **<0.0001** | **<0.0001** | **<0.0001** | 0.0501 | **<0.0001** | **<0.0001** | 0.6212 |
| | CTR-VGB – RHEB-p.P37L-VH | 0.688 | **0.0003** | **<0.0001** | **<0.0001** | **<0.0001** | **<0.0001** | **<0.0001** | **<0.0001** | **<0.0001** | **<0.0001** | **<0.0001** |
| | CTR-VGB – RHEB-p.P37L-VGB | 0.3367 | **0.003** | **<0.0001** | **<0.0001** | **0.0066** | **<0.0001** | **0.0007** | **0.0058** | 0.079 | >0.9999 | **0.0149** |
| **%RS** | CTR-VH – RHEB-p.P37L-VH | | **<0.0001** | **0.0006** | **<0.0001** | **<0.0001** | **<0.0001** | **<0.0001** | **<0.0001** | **<0.0001** | **0.0041** | >0.9999 |
| | CTR-VGB – RHEB-p.P37L-VH | | **0.0004** | **0.0008** | **<0.0001** | 0.1017 | **0.0027** | 0.0782 | **0.0078** | **0.0006** | 0.2811 | 0.7776 |
| | CTR-VGB – RHEB-p.P37L-VGB | | **0.001** | **<0.0001** | **<0.0001** | **0.0004** | **<0.0001** | 0.9121 | >0.9999 | >0.9999 | >0.9999 | 0.2372 |
| **NBR** | CTR-VH – RHEB-p.P37L-VH | | **<0.0001** | **<0.0001** | **<0.0001** | **<0.0001** | **<0.0001** | **<0.0001** | **0.0002** | **<0.0001** | **0.0019** | >0.9999 |
| | CTR-VGB – RHEB-p.P37L-VH | | **0.0002** | **<0.0001** | **<0.0001** | **<0.0001** | **<0.0001** | **<0.0001** | **<0.0001** | **<0.0001** | **<0.0001** | **<0.0001** |
| | CTR-VGB – RHEB-p.P37L-VGB | | **0.001** | **<0.0001** | **<0.0001** | >0.9999 | >0.9999 | >0.9999 | **0.006** | 0.1569 | 0.3992 | **0.0015** |
| **NIBI** | CTR-VH – RHEB-p.P37L-VH | **0.0004** | **<0.0001** | **0.0002** | **0.0003** | **0.0004** | **0.0083** | **0.0025** | **0.0153** | >0.9999 | **0.0004** | **<0.0001** |
| | CTR-VGB – RHEB-p.P37L-VH | **0.0165** | **0.0001** | **<0.0001** | **<0.0001** | **<0.0001** | | 0.0867 | **0.0028** | **0.0052** | **0.0165** | **0.0001** |
| | CTR-VGB – RHEB-p.P37L-VGB | **0.0014** | **0.0002** | 0.9663 | >0.9999 | 0.5108 | | >0.9999 | 0.2728 | >0.9999 | **0.0014** | **0.0002** |
| **$CoV^{NIBI}$** | CTR-VH – RHEB-p.P37L-VH | >0.9999 | **0.0002** | **0.0144** | 0.2102 | >0.9999 | >0.9999 | **0.0002** | >0.9999 | **0.0018** | **<0.0001** | >0.9999 |
| | CTR-VGB – RHEB-p.P37L-VH | 0.0131 | **0.0093** | >0.9999 | **<0.0001** | **0.0337** | 0.2976 | | 0.5731 | 0.08 | 0.4497 | **0.0131** |
| | CTR-VGB – RHEB-p.P37L-VGB | 0.0064 | **0.0051** | >0.9999 | 0.7631 | 0.5641 | 0.5681 | | >0.9999 | >0.9999 | >0.9999 | **0.0064** |
| **NBD** | CTR-VH – RHEB-p.P37L-VH | | **0.0018** | >0.9999 | 0.8203 | >0.9999 | >0.9999 | 0.8622 | 0.199 | **0.0289** | >0.9999 | 0.7055 |
| | CTR-VGB – RHEB-p.P37L-VH | | **0.0154** | >0.9999 | >0.9999 | >0.9999 | >0.9999 | 0.1306 | **0.0002** | 0.1921 | >0.9999 | **0.0205** |
| | CTR-VGB – RHEB-p.P37L-VGB | | **0.006** | 0.6544 | >0.9999 | **<0.0001** | **<0.0001** | **<0.0001** | **<0.0001** | **0.0178** | 0.4308 | 0.8911 |
| **NBC** | CTR-VH – RHEB-p.P37L-VH | | **0.0022** | >0.9999 | >0.9999 | >0.9999 | 0.1313 | 0.2556 | >0.9999 | **0.002** | 0.5146 | >0.9999 |
| | CTR-VGB – RHEB-p.P37L-VH | | **0.0254** | >0.9999 | 0.244 | **0.0019** | **0.0005** | **<0.0001** | **<0.0001** | **<0.0001** | **<0.0001** | **<0.0001** |
| | CTR-VGB – RHEB-p.P37L-VGB | | **0.004** | >0.9999 | 0.244 | >0.9999 | **0.0348** | **0.004** | **0.0331** | **0.0011** | >0.9999 | **0.0362** |
| **Reverberation duration** | CTR-VH – RHEB-p.P37L-VH | | **0.0055** | **0.0255** | **<0.0001** | 0.0724 | **0.0014** | **<0.0001** | **0.0002** | 0.0585 | **<0.0001** | **<0.0001** |
| | CTR-VGB – RHEB-p.P37L-VH | | **0.0116** | **0.0175** | **<0.0001** | **<0.0001** | **<0.0001** | **<0.0001** | **0.0001** | **<0.0001** | **<0.0001** | **<0.0001** |
| | CTR-VGB – RHEB-p.P37L-VGB | | **0.0046** | **0.0073** | **<0.0001** | **0.0002** | >0.9999 | >0.9999 | 0.118 | 0.1922 | **<0.0001** | **0.0315** |

**MFR = Mean firing rate, %RS = percentage random spikes, NIBI = network inter-burst interval, $CoV^{NIBI}$ = Coefficient of variance of NIBI, NBR = Network burst rate, NBD = Network burst duration, NBC = Network burst composition.**
